# Pseudogenes in plasmid genomes reveal past transitions in plasmid mobility

**DOI:** 10.1101/2023.11.08.566193

**Authors:** Dustin M. Hanke, Yiqing Wang, Tal Dagan

## Abstract

Evidence for gene non-functionalization due to mutational processes is found in genomes in the form of pseudogenes. Pseudogenes are known to be rare in prokaryote chromosomes, with the exception of lineages that underwent an extreme genome reduction (e.g., obligatory symbionts). Much less is known about the frequency of pseudogenes in prokaryotic plasmids; those are genetic elements that can transfer between cells and may encode beneficial traits for their host. Non-functionalization of plasmid-encoded genes may alter the plasmid characteristics, e.g., mobility, or their effect on the host. Analyzing 10, 832 prokaryotic genomes, we find that plasmid genomes are characterized by threefold-higher pseudogene density compared to chromosomes. The majority of plasmid pseudogenes correspond to deteriorated transposable elements. A detailed analysis of enterobacterial plasmids furthermore reveals frequent gene non-functionalization events associated with the loss of plasmid self-transmissibility. Reconstructing the evolution of closely related plasmids reveals that non-functionalization of the conjugation machinery led to the emergence of non-mobilizable plasmid types. Examples are virulence plasmids in *Escherichia* and *Salmonella*. Our study highlights non-functionalization of core plasmid mobility functions as one route for the evolution of domesticated plasmids. Pseudogenes in plasmids supply insights into past transitions in plasmid mobility that are akin to transitions in bacterial lifestyle.

## Introduction

Pseudogenes are gene copies that have been rendered non-functional due to mutations that disrupt their expression and/or function. Non-functionalization events that generate unitary pseudogenes, which lack a functional homolog in the genome are rather rare (e.g., (1)). In eukaryotes, pseudogenes frequently originate from non-functionalization of gene copies following gene duplication (2). In prokaryotes, protein family diversification is more often mediated by horizontal gene transfer (HGT) that introduces xenologs in the recipient genome, compared to gene duplication (i.e., paralogs) (3, 4). Nonetheless, acquired genes may become non-functional upon the integration into the genome (5), e.g., due to incompatibility with the transcriptional or regulatory network of the recipient organism. Gene families of transposable elements (TEs) (e.g., transposases and site-specific recombinases) are characterized by a high frequency of pseudogenes, indicating that transposition and translocation play a major role in the evolution of pseudogenes (6). Pseudogenes thus correspond to molecular fossil record of past genes; the sequence similarity between pseudogenes and their functional homolog is expected to decay due to neutral mutational processes over time (7).

The frequency of pseudogenes in prokaryotic genomes is lower compared to eukaryotes, with several exceptions that include the genomes of *Mycobacterium leprea*, *Rickettsia prowazekii* (8), as well as *Salmonella enterica* serovars Typhi and Paratyphi A (9). It has been previously suggested that non-functional DNA in prokaryotes is rapidly eliminated by deletions due to the energetic cost of gene expression (10). The prevalence of pseudogenes in prokaryotes may be furthermore associated with bacterial lifestyles. Genomes of facultative pathogens are characterized by a high pseudogene frequency compared to free-living bacteria (11, 12), which may correspond to an initial step in the evolution of genome reduction that is common in the adaptation to an obligate host-associated lifestyle (13).

Plasmids are extrachromosomal elements that play an important role in prokaryote evolution (14). The evolution of plasmids differs from that of bacterial chromosomes due to their relatively small genome size, variable copy number in the cell, and their ability to horizontally transfer between cells. Plasmids can be either self-transmissible (i.e., conjugative), mobilizable, or non-mobilizable, and this property can be inferred from their gene content. Plasmids having the complete set of genetic features required for conjugation, e.g., genes encoding the mating pair formation (MPF) machinery, are classified as self-transmissible; a plasmid is classified as mobilizable, if it encodes an *oriT* binding site or relaxase genes, but lacks the MPF machinery gene markers (e.g., (15)). Mobilizable plasmids are able to hijack the conjugation machinery of other plasmids for horizontal transfer (reviewed in (16)). Plasmids lacking an *oriT* and a relaxase gene are typically classified as non-mobilizable (e.g., (15)). Nonetheless, it has been suggested that plasmids classified as non-mobilizable with yet unidentified *oriT* might utilize relaxases of conjugative or mobilizable plasmids for transfer (17). Plasmid mobility is an evolvable property, for example, an evolution experiment of plasmid persistence in *Escherichia coli* under strong selection for the plasmid-encoded trait exemplified a loss of plasmid self-transmissibility due to a large deletion of genes required for conjugation (18). Indeed, a recent large-scale study of plasmid genomes suggested that conjugative plasmids frequently lose their self-transmissibility, with consequent changes in the plasmid genetic repertoire due to gene non-functionalization and loss (19). Hence, plasmids that harbor conjugative origins of transfer but lacking a relaxase and MPF machinery may evolved from ancestral conjugative plasmids (17).

Here we study the extent of gene non-functionalization in plasmid genomes. For that purpose, we compare the prevalence of pseudogenes between plasmids and chromosomes using a large-scale analysis of 10, 832 publicly available genomes of 738 prokaryote genera. We further study the evolutionary history of plasmid-encoded pseudogenes in 2, 441 *Enterobacteriaceae* isolates including the genera *Klebsiella*, *Escherichia*, and *Salmonella*. Comparing closely related plasmids (i.e., homologous plasmids), we further reconstruct the origin of pseudogenes in the context of transitions in plasmid mobility.

## Materials & Methods

### Genomes data and gene families

Genomes of isolates comprising plasmids were downloaded from the NCBI RefSeq database (version 8/2022). Genome assemblies containing replicons labeled other than chromosomes or plasmids were excluded from the analysis. Only genomes with a single chromosome were retained. To avoid potential inclusion of experimentally miniaturized or genomes of low quality, we excluded genomes with chromosome size below the norm of free-living bacterial organisms (<550 kb). This threshold was determined according to the genome size of *Mycoplasma genitalium* (chromosome size ca. 580 kb) that we considered as reference for the minimal chromosome size of a free-living prokaryote species (20). The curated data consisted of 10, 832 genome assemblies from 738 prokaryotic genera (Supplementary Table S1). For a detailed analysis of pseudogenes in plasmids, we used the previously established KES dataset of enterobacterial genomes (21). The KES dataset includes 1, 114 chromosomes and 3, 098 plasmids from *Escherichia*; 755 chromosomes and 2, 693 plasmids from *Klebsiella*; 572 chromosomes and 993 plasmids from *Salmonella* (all from RefSeq version 01/2021; Supplementary Table S2). Protein coding genes in the KES genomes set are clustered into 32, 623 gene families as previously described in (21). Briefly, reciprocal best hits (RBHs) of protein sequences between all replicon pairs were identified using MMseqs2 (22) (v.13.45111, with module easy-rbh applying a threshold of E-value ≤ 1 × 10^−10^). RBHs were further compared by global alignment using parasail-python (23) (v. 1.2.4, with the Needleman-Wunsch algorithm). Sequence pairs with ≥30% identical amino acids were clustered into gene families using a high-performance parallel implementation of the Markov clustering algorithm (24) (HipMCL with parameter --abc -I 2.0).

### Retrieval of pseudogenes

Pseudogenes were extracted from the RefSeq genomes according to their annotation and coordinates in the RefSeq genome feature tables. Pseudogenes in RefSeq genomes are annotated by the NCBI Prokaryotic Genome Annotation Pipeline (PGAP) (25). Briefly, PGAP utilizes sequence similarity-based approaches and *ab initio* gene prediction to annotate genomes (26). Pseudogene annotation is part of the PGAP pipeline for the annotation of protein-coding genes. Genomic loci that have sequence similarity to known genes, yet are lacking evidence of other gene features (e.g., complete open reading frame), are annotated as pseudogenes.

The quality of PGAP pseudogene annotation in KES dataset was further validated using Pseudofinder (27) (v.1.1.0, with module annotate and parameters -it 0.95 -s 0.8 -e 1e-9). The output of Pseudofinder was filtered to include only pseudogenes identified in intergenic regions. That is, pseudogenes annotated by Pseudofinder in the same locus as functional genes in the PGAP annotation were excluded. All of the PGAP annotated pseudogenes could be validated using Pseudofinder (see Supplementary text, Supplementary Fig. S1 & S2). Pseudogenes identified solely by Pseudofinder were excluded.

All KES pseudogenes were furthermore tested for programmed frameshifts (as described, e.g., for bacterial insertion sequences (ISs) IS*1*, IS*150*, and IS*911* (28)). Therefore, we searched for each pseudogene for a similar, large complete protein sequence in the KES dataset using MMseqs2 (22) (v.13.45111, with module search and parameters: --min-seq-id 0.95 -e 1.000E-09 -c 0.95 --cov-mode 2). The search result was filtered for the best alignment for each pseudogene using MMseqs2 (22) (v.13.45111, with module filterdb and parameter --extract-lines 1). The protein sequences were used as a query for sequence search against the contig where the corresponding pseudogene was found with BLAST (29) (v.2.12.0+, with module tblastn applying a threshold of E-value ≥ 1 x 10^-9^). This procedure yielded 135 (0.03%) pseudogenes that were identical to a complete protein sequence. The annotated function of those protein-coding genes corresponded mostly to hypothetical proteins (115 out of 135). Hence, we find no significant evidence for PGAP misclassification of pseudogenes due to programmed frameshifts of ISs.

### Assignment of pseudogenes into gene families

Pseudogenes were assigned into the KES gene families using their nucleotide sequences that were extracted from genome assemblies according to the genomic coordinates annotated by PGAP. Pseudogenes were aligned with the amino-acid sequences of the KES gene families using MMseqs2 (22) (v.13.45111, with module search applying a threshold of E-value ≤ 1 × 10^−10^). These MMseqs2 parameters trigger an alignment of all six translated reading frames of the pseudogene nucleotide sequence to the target amino acids sequence. Pseudogenes were assigned into the gene family that had the highest number of significantly aligned gene family members to the pseudogene. The final KES dataset encompassed 32, 623 gene families consisting of 11, 995, 860 protein coding genes and 393, 350 (96% or the total) pseudogenes in 13, 375 gene families. The annotation of gene function was extracted from the RefSeq database and gene families were assigned the most frequent annotation among their member genes. The annotation of specific gene families discussed in detail in the results was further validated using sequence search against TnCentral (30) and Pfam (31) using the most frequent sequence variant of the gene families as query. During this analysis, pseudogenes in a single relaxase/nuclease domain-containing gene family were suspected as erroneously classified as such, and were excluded from the analysis of ColE1-like plasmids. The excluded pseudogenes were annotated in the sequences of 487 plasmids. The pseudogene had a highly conserved synteny within mobilization related functions such as *mobC*, *mbeB,* and *mbeD*. In addition, homologous ORFs to *mbeA* (e.g., WP_077943844.1) were identified at the same genomic location supporting the classification of that locus as a gene rather than pseudogene.

### Inference of dN and dS distances

As a first step, the most similar gene sequence to the pseudogene was identified. For that, pseudogene nucleotide sequences were translated into amino acid sequences using transeq from EMBOSS (32, 33) (v.6.6.0.0, with parameters -frame 6 and -table 11). Translated pseudogenes were searched against the KES protein sequences using BLAST (29) (v.2.14.0+, with module blastp). The best hit was used for further comparison. A pairwise alignment between the translated pseudogene and the best matching amino acid sequence was inferred using Clustal Omega (34) (v.1.2.4, --outfmt=clu). Codon alignment was calculated using pal2nal (35) (v.14.1, with parameters -output paml and -codontable 11). The dN and dS distances were calculated using codeml from PAML (36) (v.4.9, with runmode = -2). Additionally, the maximum truncation size of pseudogenes compared to their homologous genes was determined by calculating the leading and trailing truncations between the pseudogene and the best matching homologous gene.

### Inference of unitary and non-unitary pseudogenes

The closest homologous gene of pseudogenes in the KES dataset were inferred using MMseqs2 (22) (v.13.45111, with module search applying thresholds of E-value ≤ 1 × 10^−10^ and amino acid identity ≥ 30%). Thereby, four scenarios (categories) of the homologous gene locus were examined in the following hierarchical order: (I) on the same replicon, (II) on the same replicon type, (III) on a different replicon type, and (IV) unitary pseudogene (see illustration of Fig. 1B). In the example of plasmid pseudogenes, this analysis determined homologous protein-coding genes from the same plasmid, other plasmids of the same isolate, or the chromosome of the same isolate. If no homolog was found in the same bacterial isolate, the pseudogene was classified as unitary. The same procedure was applied for chromosomal pseudogenes identifying homologous genes on the same chromosome or plasmids within the same bacterial isolate.

**Figure 1.**
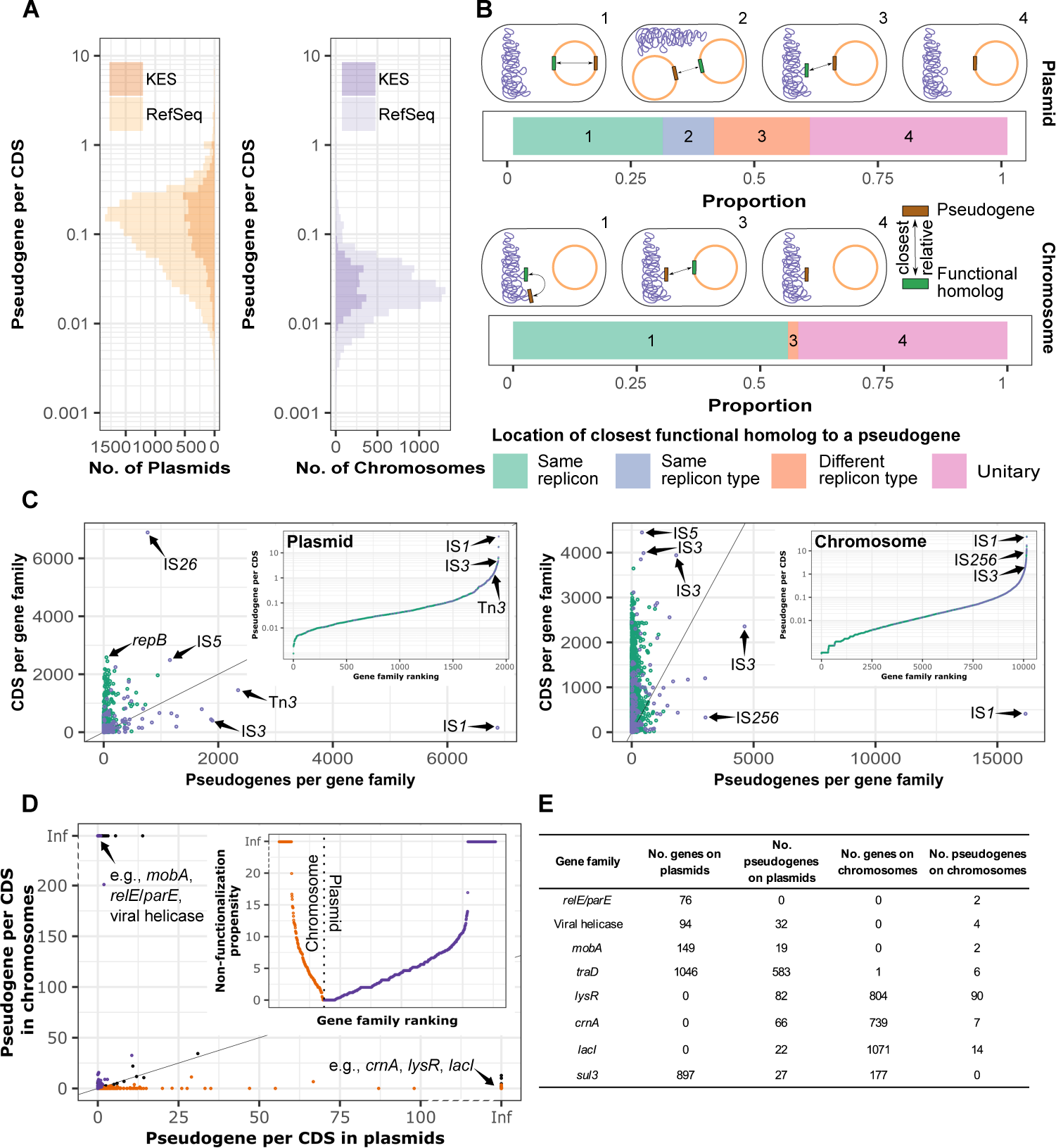
Evolution of gene non-functionalization in plasmids compared to chromosomes. **(A)** Distribution of pseudogene density (pseudogene per CDS) per plasmid and chromosome for the RefSeq and KES datasets. The graph includes plasmids and chromosomes with pseudogene content (RefSeq dataset: 21, 564 (74%) plasmids and 10, 832 (100%) chromosomes; KES dataset: 5, 573 (82%) plasmids and 2, 441 (100%) chromosomes). An increased pseudogene density in plasmids compared to chromosomes was observed also when analyzing pseudogenes detected with Pseudofinder (see Supplementary text, Supplementary Fig. S1) **(B)** Proportions of pseudogenes according to the location of the closest related homologous CDSs in plasmids and chromosomes. The median sequence similarity between pseudogenes and their homologous genes ranged between 59-87% in the five location categories (Supplementary Fig. S6). **(C)** Non-functional to functional ratio per gene family in plasmids and chromosomes. Each dot corresponds to a gene family (green). Transposase gene families are marked (purple). **(D)** Ratio of pseudogene per CDS in plasmids (x-axis) to pseudogene per CDS in chromosomes (y-axis) for 6, 396 gene families that occur on both plasmids and chromosomes. Gene functions with a significant enrichment for pseudogenes are highlighted for plasmids (orange, n=840) and chromosomes (purple, n=221) (P < 0.001, using Fisher’s exact test). Infinite (Inf) values correspond to gene families lacking pseudogenes (i.e., divisions by zero). **(E)** Example gene families that are enriched for pseudogenes. Names of listed gene families **(C, D, E)** have been validated with Pfam (31) and TnCentral (30) databases using sequence search with the most frequent sequence variant of the gene families.

### Plasmid typing and inference of closely related (homologous) plasmids

Plasmid mobility type was predicted using MOB-suite (15) (v.3.0.3). The inference of closely related plasmids was based on a new scheme of plasmid taxonomic units (PTUs) (37) and shared gene content. Plasmids were assigned to plasmid taxonomic units (PTUs) using COPLA (38) supplying taxonomic information of their hosts (Supplementary Table S3). Plasmid taxonomic units correspond to a classification scheme of closely related plasmids as inferred from average nucleotide identity (ANI) networks of whole plasmid sequences (38). Additionally, plasmids were clustered into clusters of homologous plasmids based on shared gene content. Pairwise shared gene and pseudogene content of plasmids were calculated with the Jaccard similarity index from the gene and pseudogene families. The shared gene and pseudogene content were visualized with hierarchical clustering using pheatmap (v.1.0.12). Plasmid pairs sharing ≥ 60% of their genes were filtered to construct plasmid clusters using the mcl algorithm (39) (v.12-135, parameter -- abc -I 2.0).

### Phylogenetic analysis

Plasmid and host phylogenies were constructed from complete single copy gene families for the analyzed set of plasmids or chromosomes. Sequences were aligned using MAFFT (40) (v.7.520, with parameter --maxiterate 1000 --localpair). Host phylogeny in 133 universal chromosomal genes was reconstructed using IQ-TREE 2 (41) (v.2.2.2.7) with Le and Gascuel (LG) model based on amino acid sequences (with parameter -m LG -B 1000, (42)). The host phylogeny was rooted at midpoint using iTOL (43) (v.6.8). The reconstruction of plasmid phylogenetic trees was based on nucleotide sequences of complete single-copy genes (see Supplementary Table S4 & S5) applying IQ-TREE 2 (41) (v.2.2.2.7) with a transition model (with parameter -m TIM -B 1000, (42)). The root position in the plasmid phylogenies was inferred by applying phylogenomic rooting using nucleotide sequences from plasmid gene families (44) (Supplementary Table S4 & S5). The obtained phylogenetic trees were visualized and annotated using iTOL (43) (v.6.8).

### Statistics

All statistical tests were performed using R (v.4.0.3).

## Results

### Higher density of pseudogenes in plasmids compared to chromosomes

To examine the presence of pseudogenes in plasmids and chromosomes we surveyed 10, 832 genome annotations from the Prokaryotic Genome Annotation Pipeline of NCBI’s RefSeq dataset (25). Pseudogenes were annotated in 21, 564 (74%) plasmids and 10, 832 (100%) chromosomes. Our results reveal that plasmids are characterized by a higher pseudogene density (pseudogene per CDS) compared to chromosomes (Fig. 1A), with a median of 0.070 pseudogenes per coding sequence (CDS) for plasmids and 0.024 for chromosomes (P < 0.001, using Wilcoxon test). This equates to approximately 14 CDS per pseudogene for plasmids and 42 CDS per pseudogene for chromosomes. Thus, the genomic composition of plasmids is enriched by a 3-fold higher pseudogene density compared to chromosomes.

The size difference between plasmids and chromosomes raises the question of whether replicon size (i.e., plasmid or chromosome size) may explain the different pseudogene densities. To test the impact of replicon size differences, we constructed a dataset comprising chromosomes and plasmids with similar size distributions (Supplementary Fig. S3). Comparing the pseudogene density between plasmids and chromosomes in that set showed that, also for plasmids and chromosomes of similar size, plasmids have a significantly higher pseudogene density compared to chromosomes (P < 0.001, using permutation test, median_plasmid_=0.06, median_chromosome_=0.03). Hence, the smaller plasmid genome size cannot alone explain the high pseudogene density in plasmids.

To test if the high pseudogene density in plasmids is a general phenomenon, we compared the pseudogene density between plasmids and chromosomes within different genera. Our results show that the pseudogene density was consistently higher in plasmids compared to chromosomes for all taxa, except *Shigella* (P < 0.001, using Wilcoxon test with FDR correction, Supplementary Fig. S4). The high frequency of gene non-functionalization in *Shigella* has been previously associated with adaptation of a host-associated lifestyle in this genus (11, 45). Hence, the elevated occurrence of pseudogenes in plasmids is common to most taxonomic groups.

To study plasmid gene non-functionalization in detail, we focused on plasmids residing in 2, 441 *Klebsiella*, *Escherichia*, and *Salmonella* (KES) isolates. These three enterobacterial taxa have been extensively sampled for sequencing and plasmids in these taxa include diverse plasmid types (21, 37). The density of pseudogenes in KES plasmids is significantly higher compared to chromosomes (P < 0.001, using Wilcoxon test, Fig. 1A). This reflects an observed 4-fold increase in pseudogene density in plasmids compared to chromosomes and equates to approximately 9 CDS per pseudogene in plasmids and 40 CDS per pseudogene in chromosomes. Hence, enterobacterial genomes within the KES dataset capture the trend of increased pseudogene density in plasmids compared to chromosomes that we observed within the taxonomically diverse RefSeq dataset.

Pseudogenes having a functional homolog in the isolate genome, including the chromosome and extra-chromosomal elements, may have evolved from gene non-functionalization following a gene duplication (either within the same replicon, or via translocation of duplicates between replicons). Pseudogenes that do not have a functional homolog within the same isolate may have evolved either from non-functionalization of a single-copy gene, or alternatively, non-functionalization of a horizontally acquired gene upon arrival. The comparison of pseudogenes with their functional homologs showed that most pseudogenes are truncated (289, 746, 71%). Recent studies suggest that early stop codons in many truncated pseudogenes correspond to nonsense mutations (46), or transient alleles that may be reversed under purifying selection (47). Comparing the evolutionary rates of truncated and non-truncated pseudogenes we find that truncated pseudogenes are characterized by higher dN and dS, as well as higher dN/dS compared to non-truncated pseudogenes (Supplementary Fig. S5), as is expected for DNA evolution following a relaxation of selection pressure (i.e., gene non-functionalization). Furthermore, the divergence of pseudogenes in plasmids and chromosomes compared to their closest homologous gene had similar characteristics (Supplementary Fig. S5), hence the high density of pseudogenes in plasmids cannot be explained by a bias in the genome annotation pipeline.

Pseudogenes were further classified into two groups: unitary and non-unitary pseudogenes, based on whether a functional homolog to the pseudogene was present within the same isolate. Our results show that the proportion of pseudogenes that are accompanied by a homologous gene on the same replicon is lower for plasmid pseudogenes (Fig. 1B, see also Supplementary Fig. S6). The proportion of unitary pseudogenes at the isolate level is comparable between plasmids (40%) and chromosomes (42%). Plasmid pseudogenes may furthermore have a homologous gene either in the chromosome (19%) or in another plasmid (10%) within the same isolate. Chromosomal pseudogenes having a plasmid homologous gene are rather rare (2%). Plasmid pseudogenes are thus more likely to have a homologous gene on a different replicon within the isolate genome compared to chromosomal pseudogenes. Accordingly, we conclude that gene non-functionalization in plasmids frequently entails the loss of redundant functions at the level of the whole isolate genome.

### The majority of pseudogenes in plasmids correspond to deteriorated insertion sequences

Which genes are frequently non-functionalized? In plasmids, gene families comprising transposition-related gene functions such as IS*1*, Tn*3*, and IS*3* are characterized by the highest frequency of pseudogenes (Fig. 1C). Similarly, transposases such as those encoded by IS*1*, IS*256*, and IS*3*, are frequently non-functionalized in chromosomes (Fig. 1C). Non-functional insertion sequences (ISs) accounted for a substantial portion of pseudogenes in our dataset with 88, 107 (22%) pseudogene instances. For example, the IS*1* family transposase alone comprised approximately every 20^th^ pseudogene (5.85%), with an average of 33 pseudogenes per gene. Furthermore, pseudogenes in the IS*1* family transposases are the most abundant in the data and occur in 72% of the chromosomes (Supplementary Fig. S7). Our results are thus in agreement with previous reports on frequent non-functionalization of TEs in prokaryote genomes (6) and furthermore show a similar trend in plasmid genomes.

The comparison between plasmids and chromosomes further reveals that the propensity for non-functionalization may be different depending on the replicon type. We identified 221 (0.64%) plasmid-encoded gene families that are significantly enriched for pseudogenes in the chromosome (Fig. 1D, Supplementary Table 6); of these families, 84 (38.01%) include a homologous gene in the plasmid within the same isolate (i.e., as category 3 in Fig. 1B). These families often correspond to plasmid genes, such as mobilization genes (*mobA*), toxin-antitoxin systems (*relE*/*parE*), and viral helicases (Fig. 1E). Additionally, we identified 840 (2.4%) chromosomal gene families that are significantly enriched for pseudogenes in plasmids (Fig. 1D, Supplementary Table S6); of these families, 498 (59.28%) have a chromosomal homologous gene in the same isolate. These families include genes such as LysR-type transcriptional regulators (*lysR*), LacI-related transcriptional regulators (*lacI*), and creatininase (*crnA*), that are specifically encoded on the chromosome (Fig. 1E). We hypothesize that these gene families correspond to replicon-specific genes that underwent non-functionalization upon translocation to the other replicon, rendering them essentially dead-on-arrival. Barriers for gene transfer from chromosomes to plasmids have been previously shown, for example due to dose effect (termed also dose repetition) (48). The acquisition of an extra copy of a chromosomal protein-coding gene may lead to an increased dose of the product protein, which may have a negative effect on the host fitness. For example, the presence of an additional copy of the core chromosomal chaperonin genes *groEL*/*groES* on a plasmid in *E. coli* was shown to have a negative fitness effect on the plasmid host (48). Our results suggest the presence of similar barriers for gene transfer from plasmids to chromosomes.

### Frequent pseudogenes in large mobilizable and non-mobilizable plasmids

Which plasmid types are most associated with a high frequency of pseudogenes? Initially we focus on two prominent plasmid characteristics: plasmid size and mobility class. The small plasmid types in the KES dataset mostly correspond to ColE1-like plasmids; accordingly, here we consider plasmids of <19Kbp as small (21). Our analysis shows that most of the large plasmid types (4, 659, 97%) and about half of the small plasmid types (1, 045, 54%) contain pseudogenes. The pseudogene density is highest in small non-mobilizable plasmids (small plasmid types with pseudogenes: median_small mobilizable_=0.2, median_small non-mobilizable_=0.33) compared to large plasmid types and chromosomes, but that property is tightly linked to their small size encoding only few genes. Among the large plasmids, pseudogene density is highest in mobilizable plasmids followed by non-mobilizable plasmids (P < 0.05, using pairwise Wilcoxon test with FDR correction, median_large mobilizable_=0.27, median_large non-mobilizable_=0.15, median_large conjugative_=0.1, Fig. 2A). Thus, approximately every fourth gene is non-functionalized in some mobilizable plasmids. Taken together, our results point towards an association between increased pseudogene densities in large plasmids and a lack of plasmid mobility.

**Figure 2.**
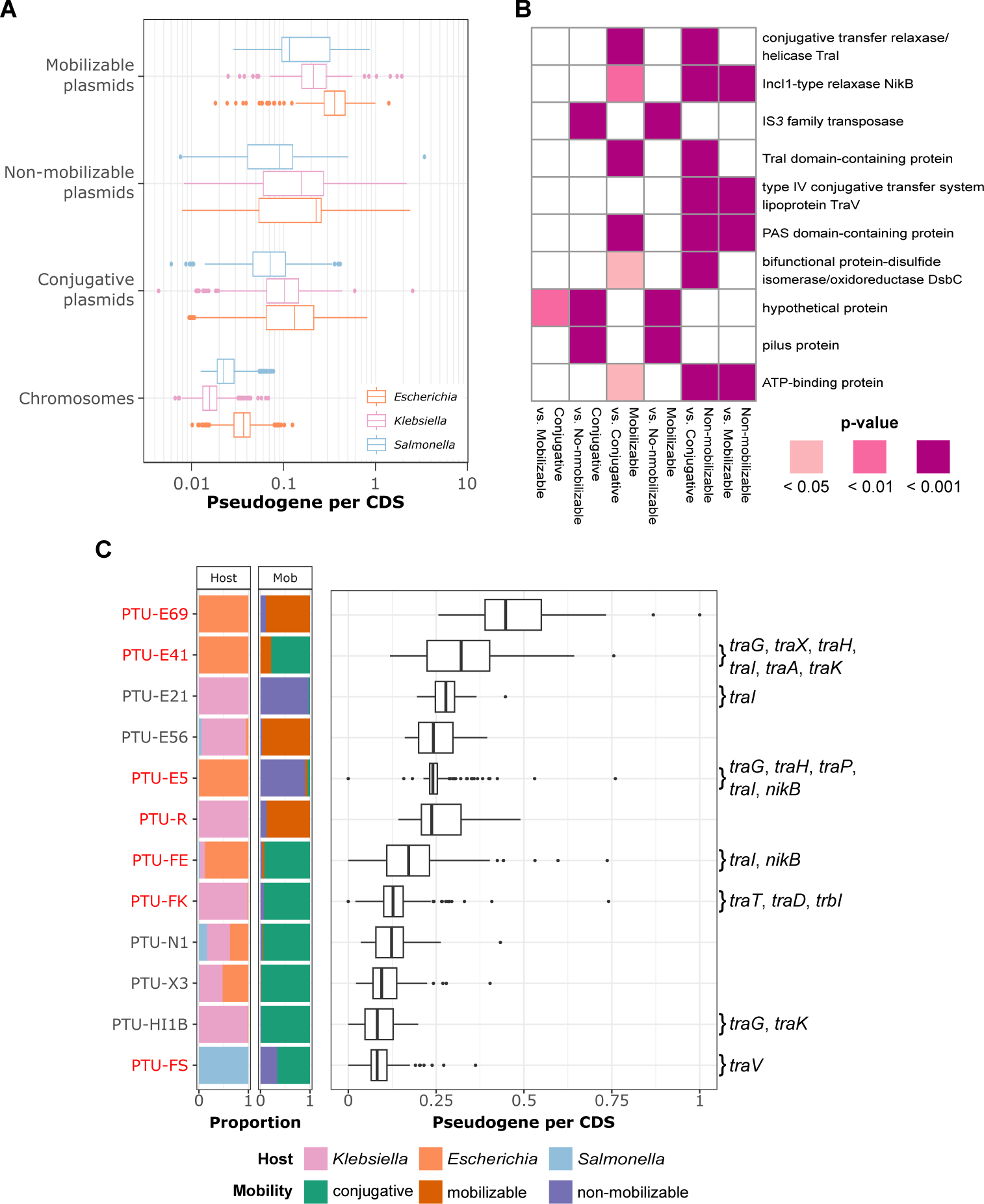
Pseudogene density and gene function in large (≥19Kb) plasmid types. **(A)** Ratio of pseudogene per CDS shown for large plasmid types (≥19Kb) and chromosomes (Replicons lacking pseudogenes were excluded. This includes 94 (3%) conjugative, 9 (1.7%) mobilizable, and 61 (5.1%) non-mobilizable plasmids). **(B)** Enrichment for pseudogenes in gene families significantly depends on the plasmid mobility type in large plasmids (P < 0.05, using Fisher’s exact test with FDR correction). The first mentioned plasmid mobility type indicates in which type the enrichment of pseudogenes for gene functions has been statistically observed. The second mentioned plasmid mobility type indicates the type from which the frequencies of pseudogenes and CDSs were used for the statistical comparison. **(C)** Distribution of pseudogene per CDS ratio for plasmid taxonomic units (PTUs) of large plasmid type (only frequent (n_PTU_≥30) PTUs are shown). Stacked bar plots show proportions of plasmid mobility type and the host taxonomy per PTU. Labels besides the boxplot report PTU-specific non-functionalized gene families corresponding to transfer-related (*tra*) functions (P < 0.05, using Fisher’s exact test with FDR correction). PTUs highlighted in red are characterized by multiple plasmid mobility types (where the main plasmid mobility type < 95%).

Plasmids in the different plasmid size and mobility categories are typically different in their composition. We therefor compared the propensity of gene non-functionalization among the plasmid size and mobility groups. Only seven gene families were enriched for pseudogenes in the small plasmid types including antibiotic resistance genes (Supplementary Fig. S8, Supplementary Table S7). A total of 488 gene families were enriched for pseudogenes in at least one of the large plasmid mobility classes (Fig. 2B). The comparison between large plasmids in different mobility classes revealed a significant enrichment for pseudogenes in gene families encoding for plasmid transfer mechanisms including *traI*, *nikB*, and *traV* (Fig. 2B, Supplementary Table S8). Consequently, we hypothesized that the origin of these pseudogenes is gene non-functionalization following loss of self-transmissibility of conjugative plasmids.

To gain further understanding of gene non-functionalization events in the context of large plasmid types, we compared the pseudogene density among closely related plasmids as inferred from plasmid taxonomic units (PTUs) (37). PTUs that comprise a high pseudogene density typically include a high proportion of mobilizable and non-mobilizable plasmid types (Fig. 2C). Gene deletions and changes in genetic repertoire of plasmids may lead to a transition in plasmid mobility, which may lead to PTU divergence and the evolution of novel PTUs in the long-term (17, 19). Indeed, here we observe several PTUs where transfer-related gene families frequently occur as pseudogenes; these PTUs are highlighted as putative plasmid types where transitions in plasmid mobility may have occurred (red labeled PTUs in Fig. 2C). Transfer-related pseudogenes thus correspond to molecular fossils of lost mobility mechanisms in plasmid genomes.

### Vertical inheritance of pseudogenes in evolutionary-related plasmid clusters

Pseudogenes observed within the same gene family may have originated either from multiple independent gene non-functionalization events, or alternatively, from few non-functionalization events in ancestral plasmids followed by vertical inheritance during plasmid diversification. If vertical inheritance of pseudogenes is common in plasmid evolution, then the number of observed pseudogenes supplies an overestimation for the frequency of gene non-functionalization events. To evaluate the role of vertical inheritance in pseudogene evolution, we compared shared gene and pseudogene content among closely related plasmids. For that purpose, we clustered all plasmids into clusters of homologous plasmids based on shared gene content. We obtained 353 clusters comprising 85% large plasmids and 206 clusters comprising 89% of the small plasmids; these clusters largely correspond to the PTUs (see Supplementary Fig. S9). A visual inspection of the shared genes matrices suggests that pseudogenes tend to be shared among plasmids that also share genes (i.e., clusters of closely related plasmid) (Fig. 3AB). Indeed, the proportion of shared genes and shared pseudogenes between plasmid pairs are positively correlated (r_s_ = 0.47, P < 0.001, using Spearman’s correlation).

The observation of shared pseudogenes in plasmids may be due to ancestral gene non-functionalization and vertical inheritance (i.e., retention) of non-functional DNA. An alternative explanation for shared pseudogene content is independent gene non-functionalization events in a narrow range of gene families. The strongest patterns of shared pseudogenes in our dataset are mostly restricted to closely related plasmid clusters. A total of 874 (10.1%) gene families comprising pseudogenes are plasmid cluster-specific (P < 0.05, using Fisher’s exact test with FDR correction). Hence, shared pseudogene content between plasmids clusters, is expected to correspond to non-functionalization events in diverse gene families. A total of 356 (4.4%) gene families correspond to frequent, non-functional transposase-related functions such as IS*1*, Tn*3*, and IS*26*. Additionally, gene family size and the frequency of non cluster-specific pseudogenes are significantly correlated (r_s_ = 0.58, P < 0.001, using Spearman’s correlation). Shared pseudogene content between plasmid clusters thus corresponds either to horizontal gene transfer or commonly non-functionalized gene families.

### Non-functionalization following segmental deletion is a signature for the loss of self-transmissibility

To gain a better understanding on the evolutionary events at the basis of pseudogene vertical inheritance, we reconstructed the evolution of closely related plasmid clusters. For that purpose, we selected two clusters of *Salmonella* plasmids that share a high proportion of their genes and correspond mostly to PTU-FS (Supplementary Table S3, Supplementary Fig. S9). The shared gene content within the selected clusters FS_A_ and FS_B_ is 92% and 86%, respectively, with an average of 50% shared genes among plasmids in both clusters (Fig. 3C). Plasmids of cluster FS_A_ are homologous to plasmid pSLT (NC_003277.2) of *S. enterica* serovar Typhimurium and plasmids of cluster FS_B_ are homologous to plasmid pSENV (NZ_CP063701.1) of *S. enterica* subsp. *enterica* serovar Enteritidis. Cluster FS_A_ comprises only 83 conjugative plasmids and cluster FS_B_ includes mostly (46, 92%) non-mobilizable plasmids, in addition to four conjugative plasmids. The average shared pseudogene content among plasmids within the clusters is 67% or 65% for clusters FS_A_ and FS_B_, respectively. The shared pseudogene content among plasmids from different clusters was only 9%, on average, thus plasmids in the two clusters are diverged in their pseudogene content. The high proportion of shared gene (and pseudogene) content among plasmids in the two PTU-FS clusters supports the notion that these plasmids are of common ancestry, i.e., they are homologs, albeit with alterations in their mobility class.

**Figure 3.**
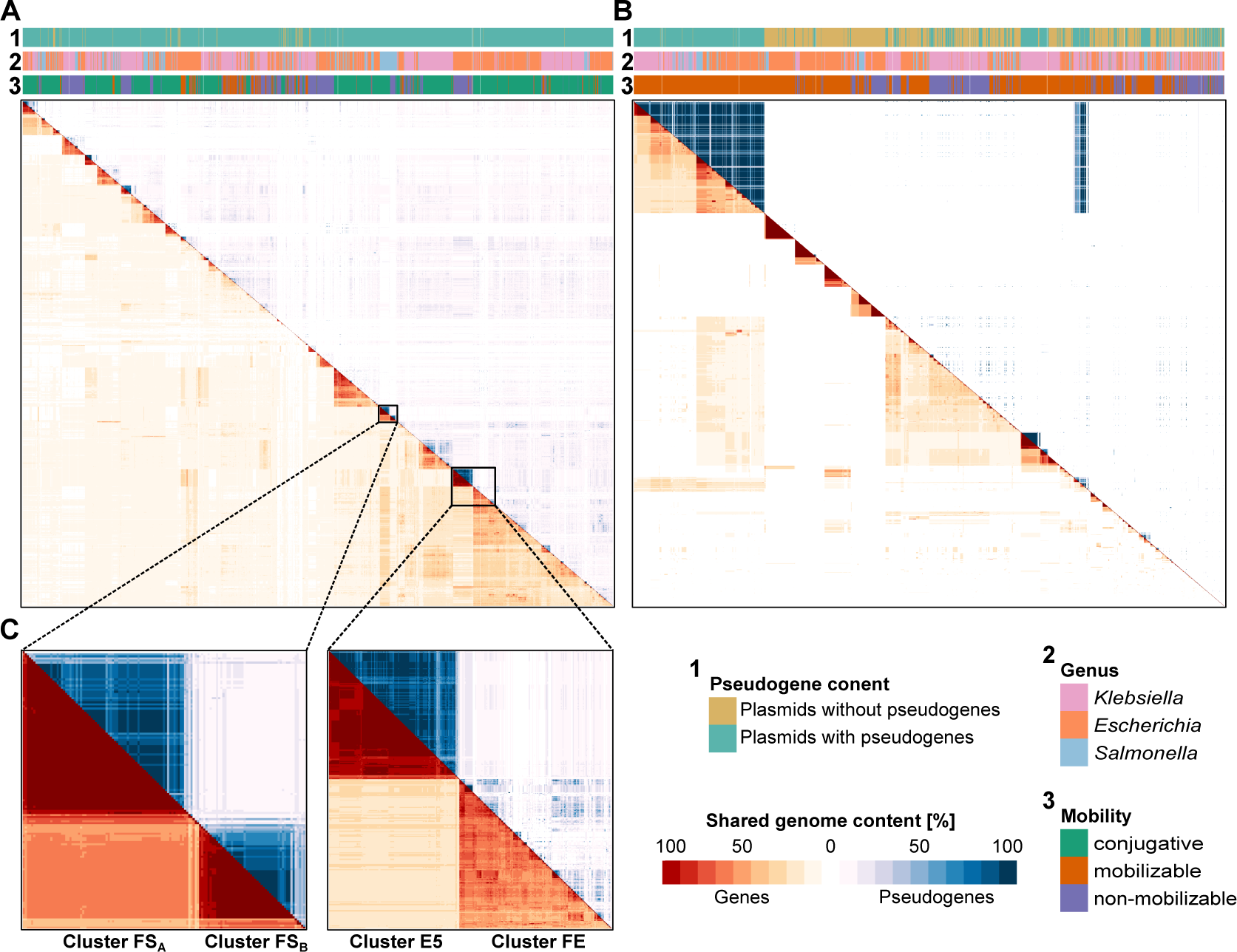
Shared gene and pseudogene content are similar in enterobacterial plasmids. **(A, B)** Pairwise shared gene (red) and pseudogene (blue) matrices of large plasmid types **(A)** and small plasmid types **(B)** among the KES dataset. The matrices were sorted using hierarchical clustering of shared gene content. **(C)** Shared genes and pseudogenes of adjacent plasmid clusters (FS_A_, FS_B_, FE, and E5) in the hierarchically clustered shared gene content matrix. Labeled plasmid clusters are based on the Markov Cluster Algorithm (MCL) using pairs of plasmids sharing ≥ 60% gene content (see methods).

To examine the evolution of PTU-FS plasmids, we reconstructed the plasmid phylogeny based on eleven single-copy gene families present in 122 (92%) plasmids. The phylogeny reveals a clear split between plasmid clusters FS_A_ and FS_B_ (Fig. 4A). To reconstruct the ancestral mobility state of PTU-FS plasmids, we inferred the root of PTU-FS using a phylogenomic rooting approach (44). The phylogenetic inference revealed a root neighborhood in the deepest split between plasmid clusters FS_A_ and FS_B_ that includes branches leading to several conjugating plasmids of cluster FS_A_ (Fig. 4A, see also Supplementary Fig. S10). This root position implies that the ancestor of PTU-FS plasmids was conjugative and the non-mobilizable plasmids in cluster FS_B_ are derived plasmids (Fig. 4A). Notably, the plasmid genome size differs significantly between the non-mobilizable cluster FS_B_ plasmids and the conjugative FS_A_ plasmids, with the non-mobilizable plasmids having a smaller genome size (P < 0.001, using Wilcoxon test, median_non-mobilizable_=59, 372 bp, median_conjugative_=93, 865 bp). Further comparison of the gene content among the PTU-FS reveals a high conservation of plasmid gene order, except for the absence of transfer (*tra*) genes in the non-mobilizable FS_B_ plasmids (Fig. 5A, Supplementary Fig. S11A).

**Figure 4.**
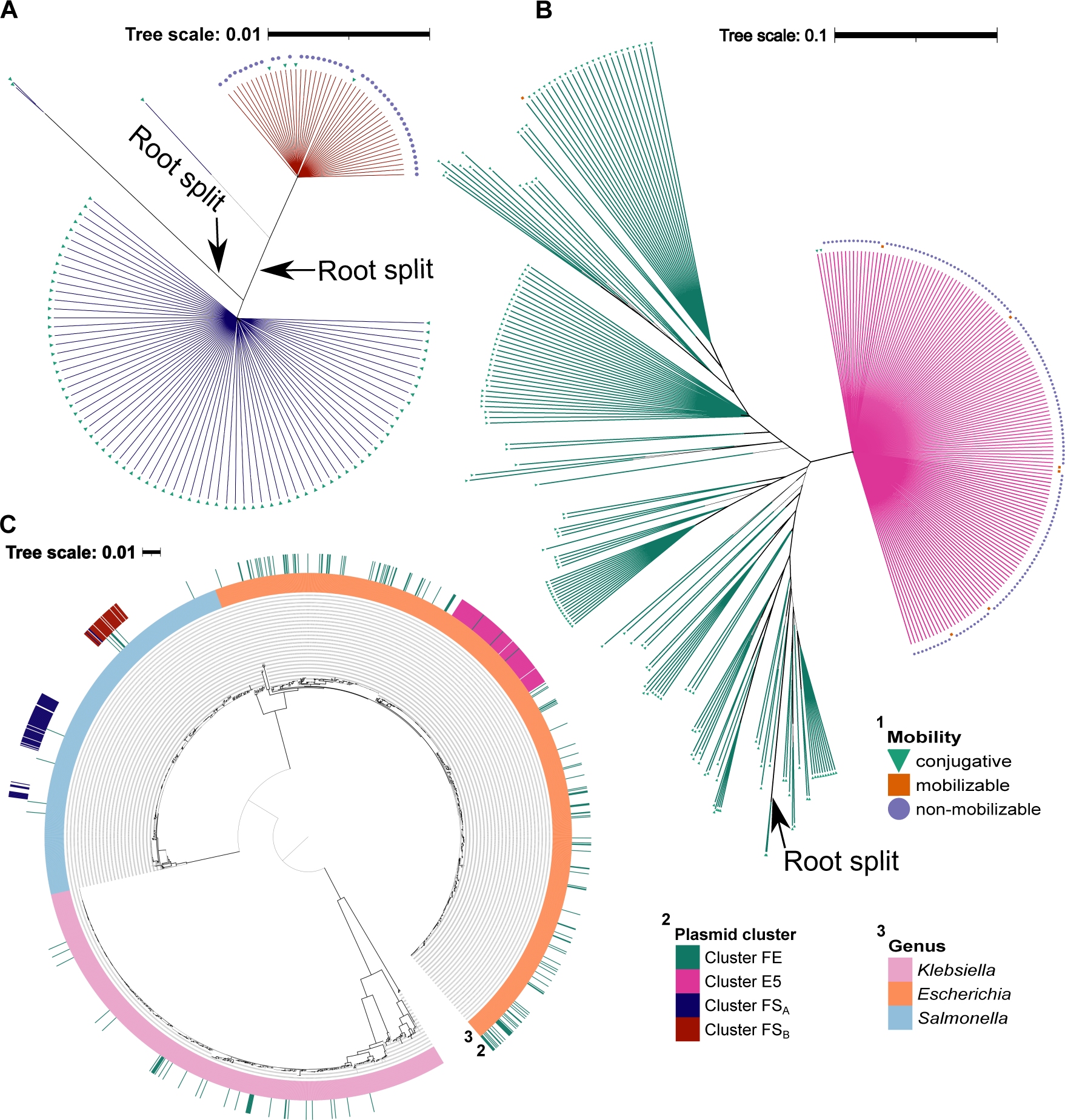
Phylogenetic relationship of self-transmissible and non-mobilizable plasmid clusters and their hosts. **(A, B)** Phylogenetic maximum-likelihood trees including 11 single-copy genes of plasmid clusters FS_A_ and FS_B_ (PTU-FS) and 10 single-copy genes of plasmid clusters FE (PTU-FE) and E5 (PTU-E5) (see Supplementary Table S4 & S5). Three plasmids (ca. 2%) and 40 plasmids (ca. 11%) were excluded from the phylogenetic reconstruction of **(A)** and **(B)**, respectively, to increase the number of complete single-copy gene families and hence the robustness of the phylogenetic inference. Arrows indicate the inferred root positions resulting from root neighborhood inference (see supporting gene trees in Supplementary Fig. S10). The operational taxonomic unit symbols correspond to predicted plasmid mobility type (legend 1). The reconstructed tree branches are colored black (note the short branch length in nearly polytomic ancestral node). The plasmid clusters (PTUs) are shown by the colored lines extending from the tree branches (legend 2). **(C)** Host phylogenetic tree based on 133 chromosomal single-copy genes. The tree was rooted using midpoint rooting. The outer color strips of the phylogenetic host tree show the cluster affiliation of plasmids residing in the hosts (legend 2) and the host genera (legend 3).

**Figure 5.**
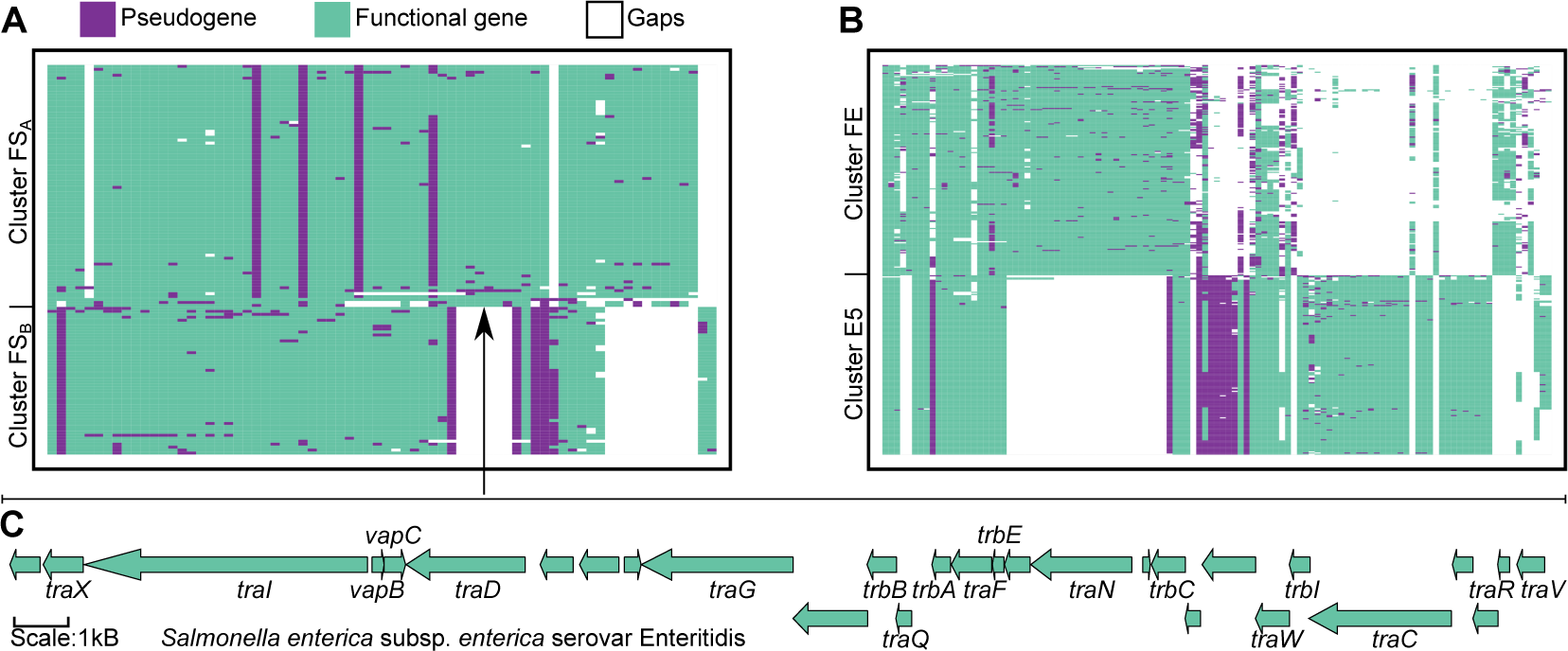
Pseudogene and gene family Presence Absence Pattern (PAP) of self-transmissible and non-mobilizable plasmid clusters. PAP of plasmid clusters FS_A_/FS_B_ **(A)** and FE/E5 **(B)**. The PAPs are sorted using hierarchical clustering of the shared gene family content. **(C)** Gene content and order in the segmental deletion inferred for the ancestor of FS_B_ (non-mobilizable) plasmids (pointed by an arrow). The inference was based on the transfer-related gene neighborhood in cluster FS_A_ (conjugative) plasmids (see also Supplementary Fig. S12).

The loss of the conjugation machinery genes may have followed two alternative scenarios: gradual non-functionalization and gene loss, or a large-scale deletion followed by non-functionalization of functionally-related genes. Since the absent genes are consecutively arranged in the ancestral plasmid (Fig. 5C), we infer that a large segmental deletion of transfer-related genes occurred in the ancestor of cluster FS_B_ non-mobilizable plasmids (Supplementary Fig. S12). The transfer-related genes *traV* and *FinO* (a repressor of conjugative transfer in F-like plasmids) correspond to the boundaries of the deleted transfer-related gene arrangement in the ancestor and are found as pseudogenes in derived plasmids. Notably, the *Salmonella* host phylogeny shows that plasmid clusters FS_A_ and FS_B_ reside in distinct groups of *Salmonella* isolates where hosts of the conjugative FS_A_ plasmids appear as more distantly related in comparison to hosts of the non-mobilizable FS_B_ plasmids (Fig. 4C, Supplementary Fig. S13). Taken together, our results reveal a transition in plasmid mobility from self-transmissible to non-mobilizable plasmids within *Salmonella* PTU-FS plasmids due to large segmental deletion followed by gene non-functionalization events.

To examine further scenarios for gene non-functionalization in plasmid genomes, we investigated two additional adjacent plasmid clusters in the shared gene matrix where the pattern of gene and pseudogene sharing indicates that they are closely related (Fig. 3C). The plasmids in these clusters correspond to the evolutionary related PTUs FE and E5 (19). Most (167, 87%) of the plasmids in cluster FE were assigned to the conjugative PTU-FE, with the remaining plasmids being unassigned (Supplementary Table S3, Supplementary Fig. S9). All 164 plasmids in cluster E5 were classified into the non-mobilizable PTU-E5 (Supplementary Table S3, Supplementary Fig. S9), which correspond to homologs of pO157 in *E. coli* strain O157:H7 and are known to be non-transmissible (17, 49). Plasmids in cluster FE share, on average, 53% of their genes and 12% of their pseudogenes. In comparison, gene sharing among plasmids in cluster E5 shows a higher shared content with 90% of the genes and 80% of the pseudogenes shared, on average. Plasmids in FE and E5 clusters share, on average, 18% gene content and 5% pseudogene content. The shared gene content among plasmids in the two clusters reflects evolutionary relationships where the different plasmid mobility types may be explained by the differential gene content.

To examine the evolutionary history of plasmids in clusters FE and E5, we reconstructed the plasmid backbone phylogeny using ten single-copy gene families present in 316 (89%) plasmids. The phylogeny revealed a clear split between the plasmid clusters (Fig. 4B). The topology of cluster E5 plasmids reveal two main plasmid lineages of closely related plasmids. In contrast, the phyletic pattern of cluster FE conjugative plasmids revealed a high divergence among the plasmids (Fig. 4B). The inferred root position is located in a branch leading to two conjugative cluster FE plasmids (Supplementary Fig. S10). Notably, two plasmids in cluster E5 encode a partial repertoire of the conjugation machinery and they were accordingly predicted as conjugative; these plasmids branch closely to the split with cluster FE plasmids (Supplementary Fig. S14) and their syntenic gene order reveals differential loss of transfer-related genes (Supplementary Fig. S14). Comparing the sequences of the conjugative plasmids of cluster E5 to their closest non-mobilizable neighbor plasmid revealed evidence for loss of plasmid mobility genes and shared genomic arrangements in the non-mobilizable cluster E5 plasmids (Fig. 5B, Supplementary Fig. S12B). The phylogenetic reconstruction suggests that the ancestral plasmid of cluster E5 plasmids (or PTU-E5) was conjugative and the plasmid mobility was lost due to a segmental deletion of the *tra* genes (Fig. 4B, Supplementary Fig. S14). Plasmids in cluster E5 share 80% of their pseudogenes, including transposases such as IS*3* (n=508), IS*1* (n=336), and IS*91* (n=320), as well as transfer-related gene families *traI* (n=164) and *nikB* (n=164) that are non-functionalized in 164 (98%) of the cluster E5 plasmids. Plasmids in cluster FE reside in a diverse set of hosts including *Klebsiella*, *Escherichia*, and *Salmonella*. In contrary, plasmids in cluster E5 are only hosted by closely related *Escherichia* isolates including strain O157:H7 (Fig. 4C); these plasmids are homologs of plasmid pO157 that is non-mobilizable (e.g., NZ_CP017435.1). Note that the hosts of cluster E5 plasmids are diverged, i.e., they do not correspond to isogenic strains (Supplementary Fig. S13). Hence the topology of cluster E5 phylogeny likely corresponds to within-host evolution via vertical inheritance. Notably, cluster E5 plasmids are characterized by significantly larger plasmid genome size, but a smaller number of CDSs compared to the conjugative cluster FE plasmids, which is well explained by the frequent pseudogenes in cluster E5 plasmids (P < 0.001, using Wilcoxon test, median_Cluster E5_=78 CDS, 19 pseudogenes, 93, 179 bp; median_Cluster FE=_86 CDS, 9 pseudogenes, 74, 922.5 bp). Hence pseudogenes may be retained as non-coding DNA in the evolution of the non-mobilizable plasmids after the loss of self-transmissibility. Our results thus reveal that plasmid divergence events may give rise to new integral replicons within their host lineage.

## Discussion

Gene non-functionalization is considered a common event following gene duplication and gene acquisition via lateral gene transfer. Previous studies suggested that bacterial genomes are often devoid of pseudogenes due to purifying selection (8). This implies that pseudogenes in bacterial genomes should be considered as ‘garbage DNA’, that is, non-functional DNA that may have an effect on the organism fitness, rather than ‘Junk DNA’ (see (50) for definitions). Our results show that pseudogenes are found in bacterial genomes and their density is highest in plasmids. We identify two main processes in plasmid evolution that are accompanied by frequent gene non-functionalization: proliferation of mobile genetic elements (MGEs) and loss of plasmid mobility (i.e., reductive plasmid evolution).

Mobile genetic elements, including transposons and integrons, are known to translocate or proliferate within and between genomes (51), thus, facilitating gene translocation and transfer, as well as genomic rearrangements (reviewed in (52)). Specifically in conjugative plasmids, ISs are known to mediate and facilitate the transfer of antibiotic resistance genes (53). At the same time, the integration of MGEs in bacterial genomes may lead to gene non-functionalization due to disruption of open reading frames at the insertion locus. Examples are previous reports on IS-mediated inactivation of the arginine biosynthesis pathway in the tsetse fly symbiont *Sodalis glossinidius* (54) and an inactivation of the plasmid-encoded nitrogen fixation genes (*nif*) in *Bradyrhizobium* isolates (55). The high frequency of MGE pseudogenes in plasmids reported here conforms previous studies highlighting the role of failed transposition events in the generation of pseudogenes in bacterial genomes (e.g., (6)). We observed that most pseudogenes are due to gene truncations and that pseudogenes in plasmids are more deteriorated. Indeed, site-specific recombination of transposable elements generates genomic hotspots for their insertion, where repeated insertions at the same locus may lead to gene truncations (reviewed in (56)). That being said, recent studies suggest that many pseudogenes are transcribed, and sometime even translated (57). Pseudogenes of MGE origin may thus correspond to proto-genes and a source for genetic innovation during MGE diversification (47). This suggestion is in line with recent studies that report pseudogene resurrection under strong selective conditions for the lost gene function. Examples include the resurrection of iron uptake in *E. coli* (58) and the resurrection of CO_2_ assimilation in *Brucella* spp. (59).

Our analysis furthermore reveals that the propensity for specific gene non-functionalization depends on the replicon type, i.e., plasmid or chromosome. This observation may be explained by selection against the presence of essential (chromosomal) genes in plasmids, e.g., due to unreliable plasmid inheritance (60) or deleterious effects of gene duplication (48). Thus, the higher pseudogene density in plasmids compared to chromosomes may be explained by the propensity of plasmid gene content to include accessory and dispensable genes. We note however, that the classification of genes into core and accessory may vary depending on the bacterial taxon, and that has an effect on the evolution of plasmid gene content in different bacterial taxa (e.g., in the context of antibiotic resistance plasmids; (61)). We hypothesize that the horizontal transfer (or translocation) of chromosomal genes into plasmids (and *vice versa*) is bound to yield effectively ‘dead-on-arrival’ pseudogenes.

Inferring rooted plasmid phylogenies that comprise related non-mobilizable and conjugative plasmids enabled us to infer the ancestral and derived plasmid variants in plasmid evolution. According to our inference, non-functionalized copies of transfer-related genes in large non-mobilizable and mobilizable plasmids are molecular fossils underlining a self-transmissible plasmid origin. The loss of plasmid self-transmissibility may lead to plasmid ‘domestication’ in the host lineage (or plasmid ‘fixation’ in the host pangenome). Domesticated plasmids are characterized by a stable vertical inheritance within the lineage that can be considered as ancestral states of chromids (62). Examples for the evolution of chromids from essential plasmids are found in plant-associated bacteria such as *Sinorhizobium meliloti* harboring a chromid, pSYMB, that encodes essential genes (63). Nonetheless, evolution of domesticated plasmids following the acquisition of essential genes was so far not considered in *Enterobacteriaceae* (48, 60). Previous studies reported a stable inheritance of plasmids pO157 in *Escherichia* and pSLT in *Salmonella* (64–66). Although, pSLT is self-transmissible, it was shown that most *Salmonella* virulence plasmids are rather vertically inherited except for pSENV of *S. enterica* subsp. *enterica* serovar Enteritidis (67). Plasmids of *S. enteritidis* bear incomplete *tra* regions (68) and virulence plasmids of serovars Enteritidis and Typhimurium are not mobilizable by other F-like plasmids even in the presence of an *oriT* region (69). The results of our analysis support the view that pSENV plasmid homologs were acquired by horizontal transfer subsequent to a segmental deletion of transfer-related gene (the evolution of serovar specific virulence plasmids reviewed in (70)). Furthermore, the phylogeny of isolates hosting pO157- and pSENV-like plasmids indicates that these plasmids were fixed in those lineages following the loss of self-transmissibility (Fig. 4C), hence they are better considered as domesticated plasmids (see also (71)).

The pseudogene repertoire and order in large non-mobilizable plasmids indicate that segmental deletions within the conjugation machinery play a major role in the initial steps of transitions in plasmid mobility. Such events may be caused, for example, by a transposon-mediated segmental deletion. A similar event was observed in an experimental evolution study of plasmid pKP33 adaptation to a naïve *E. coli* host under selection for the plasmid-encoded antibiotic resistance (18). Such a change in the plasmid genomic composition that leads to the loss of self-transmissibility is likely to trigger a ‘domino effect’ of sequential non-functionalization events of other transfer-related genes. Similar dynamics of gene non-functionalization have been previously suggested for the evolution of pseudogene content in obligatory symbionts (72). The loss of self-transmissibility may be deleterious for plasmids due to the inevitable restriction in the host range. Nonetheless, plasmids encoding a trait that is essential for the host will be maintained in the population, also if their inheritance is unstable due to selection acting at the level of host (e.g., see (73)). Another commonality in the evolution of plasmids and facultative pathogens is the interplay of MGE proliferation and pseudogene accumulation that is associated with genome miniaturization. Examples for pathogens having highly reduced (functional) genome size following massive gene decay are *Mycobacterium leprae*, *Yersinia pestis*, *Shigella flexneri*, *Rickettsia prowazekii*, *Salmonella enterica* serovars Paratyphi A, and Typhi (11, 74–78). Genome miniaturization following gene non-functionalization and loss has been furthermore described in the evolution of obligatory symbionts in heritable symbioses where deleterious mutations can be fixed by genetic drift (reviewed in (79)). Similarly to plasmids, frequent gene non-functionalization in the evolution of symbionts can lead to host range restriction that is accompanied by a shift from facultative to obligatory symbiosis. Co-adaptation plasmids that lost their mobility and their host can lead to the evolution of stable plasmid inheritance (e.g., as observed in experimental plasmid evolution studies (18, 76, 77)) and eventually plasmid domestication. Reductive genome evolution of plasmids is thus akin to the evolution of obligatory symbionts and the mechanisms involved in these processes – transposable element proliferation and gradual gene non-functionalization leading to shifts in the plasmid lifestyle – are likely similar.

## Data availability

The data and accession numbers (RefSeq database) are provided in electronic supplementary material.

## Acknowledgements

We thank Nils Hülter, Devani Picazo, Fabian Nies, Lisa Hartmann, Johannes Effe, Ishan Bhatt, and Mario Santer for critical comments on the manuscript. We thank Fenna Stücker for graphical abstract illustration. This research was supported in part through high-performance computing resources available at the Kiel University Computing Centre.

## Author contributions

D.M.H. and T.D. conceived the study. D.M.H. designed and performed the data analysis and visualizations. Y.W. prepared and provided complementary data. D.M.H. and T.D. interpreted the results and wrote the manuscript with comments and additions from Y.W.

