## Supplementary material for "Pseudogenes in plasmid genomes reveal past transitions in plasmid mobility"

### Supplementary text

#### Comparison of Pseudofinder and PGAP pseudogene detection results

To compare pseudogene candidates of PGAP and Pseudofinder (1) we calculated the overlap of pseudogene loci detected by both tools. Applying Pseudofinder to our dataset yielded 884,320 putative pseudogenes. A total of 407,657 (99.9%) of the PGAP pseudogenes were included in the Pseudofinder output (Supplementary Fig. S1). The remaining 437 (0.1%) PGAP pseudogenes putatively only identified by PGAP were also included in Pseudofinder's output, however, these loci were extended in Pseudofinder's output as this tool tends to join fragmented CDSs into a locus with PGAP pseudogenes if downstream and upstream CDS fragments located to the PGAP pseudogene share BLAST hits to the same gene (see more details under 'Fragmentated genes' in (1)). In other words, in case of CDS fragmentation, Pseudofinder may report a single pseudogene locus that includes two pseudogenes. Additional 476,226 (53.85%) pseudogenes in Pseudofinder's output were not included in the PGAP output. Taken together, the results from Pseudofinder supply a validation for PGAP pseudogene annotations. In our study, we included only pseudogenes identified by both tools.

Notwithstanding the large number of pseudogenes detected solely by Pseudofinder, our conclusion of an increased pseudogene density in plasmids compared to chromosomes still holds when analyzing pseudogenes identified by that tool. Repeating the comparison between plasmids and chromosomes using the full set of Pseudofinder pseudogenes confirmed a higher pseudogene per CDS ratio in plasmids ($median_{Pseudofinder}$=0.25) compared to chromosomes ($median_{Pseudofinder}$=0.06) ($P < 0.001$, using Wilcoxon test, Supplementary Fig. S1). Thus, our conclusion of an increased pseudogene density in plasmids compared to chromosomes is independent of the pseudogene detection tool.

To further examine the properties of pseudogenes solely identified by Pseudofinder in KES dataset, we compared their sequence length to pseudogenes detected by both tools (termed in the following 'PGAP Pseudogenes') (Supplementary Fig. S1). Our results show that Pseudofinder pseudogene sequences were significantly shorter than PGAP pseudogenes ($P < 0.001$, using Wilcoxon test, $median_{PGAP=}$596 nt, $median_{Pseudofinder}$=156 nt). Furthermore, the length of PGAP pseudogenes is closer to the median length of CDSs ($median_{KES\ CDS}$=807 nt) from the KES dataset compared to Pseudofinder pseudogenes (Supplementary Fig. S1). Together, these differences suggest that pseudogenes detected only by Pseudofinder correspond to fragmented pseudogenes.

An important aspect of our study is the inference of former gene function (i.e., the pseudogene functional homologs). Consequently, we compared the alignment quality of PGAP and Pseudofinder pseudogenes to their closest homologous gene sequences. The closest gene were searched with MMSeqs2 (2) (v.13.45111, with module search and parameters: --min-seq-id 0.95 -e 1.000E-09) and the best hit of the search was examined for each pseudogene (with MMSeqs2 module filterdb and parameter –extract-lines 1). Note that query pseudogenes are translated into all possible six reading frames and are searched against the amino acids sequences of the gene families. Comparing the alignments of PGAP and Pseudofinder pseudogenes to their closest homologous gene, we observed a shorter alignment length ($median_{Pseudofinder}$=156 aa, $median_{PGAP}$=474 aa) and a higher E-value ($median_{Pseudofinder}$= 1.042e-24, $median_{PGAP}$=1.885e-88) in Pseudofinder alignments compared to PGAP alignments (P < 0.001, using Wilcoxon test, Supplementary Fig. S2). Furthermore, the PGAP pseudogene alignments were characterized by similar query and target coverage compared to Pseudofinder pseudogene alignments (Supplementary Fig. S2). An unequal coverage of the query and target sequences may indicate a low sequence similarity in a global alignment of the total gene length (i.e., in contrast to the local alignment). To test this possibility, we inferred global alignments of pseudogenes to their best hit homologous gene sequence. For that, the pseudogene nucleotide sequence was translated in the reading frame that yielded the best aligning ORF to their homologous gene. Comparing the results of global alignments between pseudogenes and their homologous gene, we observed a lower sequence similarity of Pseudofinder pseudogenes to their homologs compared to PGAP pseudogenes (P < 0.001, using Wilcoxon test, $median_{Pseudofinder}$= 73%, $median_{PGAP}$=81%, Supplementary Fig. S2). Hence, in addition to being more fragmented, the pseudogenes identified only by Pseudofinder are also characterized by a lower sequence similarity to their homologous genes. Including in our analysis only pseudogenes identified by both tools thus implies that pseudogenes with highly ameliorated sequence similarity are excluded from our analysis.

### Supplementary Tables

**Supplementary Table S1. Genome Assemblies of the RefSeq dataset.**
See supplied Excel file SuppTables.

**Supplementary Table S2. Genome Assemblies of the KES dataset.**
See supplied Excel file SuppTables.

**Supplementary Table S3. Plasmid clusters and plasmid taxonomic units (PTUs) of plasmids in the KES dataset.**
See supplied Excel file SuppTables.

**Supplementary Table S4. Statistical report of the phylogenomic rooting result for plasmid cluster $FS_A$ and $FS_B$ (PTU-FS).**
See supplied Excel file SuppTables.

**Supplementary Table S5. Statistical report of the phylogenomic rooting result for plasmid cluster FE (PTU-FE) and E5 (PTU-E5).**
See supplied Excel file SuppTables.

**Supplementary Table S6. Non-functionalization propensities of gene families being either located on the chromosome or plasmid.**
See supplied Excel file SuppTables.

**Supplementary Table S7. Non-functionalization enrichments of gene families on large plasmid types.**
See supplied Excel file SuppTables. Values in the excel file correspond to negative logarithm of p-values to the base 10 ($-\log_{10}$) from one-sided Fisher's exact test with FDR correction.

**Supplementary Table S8. Non-functionalization enrichments of gene families on small plasmid types.**
See supplied Excel file SuppTables. Values in the excel file correspond to negative logarithm of p-values to the base 10 ($-\log_{10}$) from one-sided Fisher's exact test with FDR correction.

### Supplementary Figures

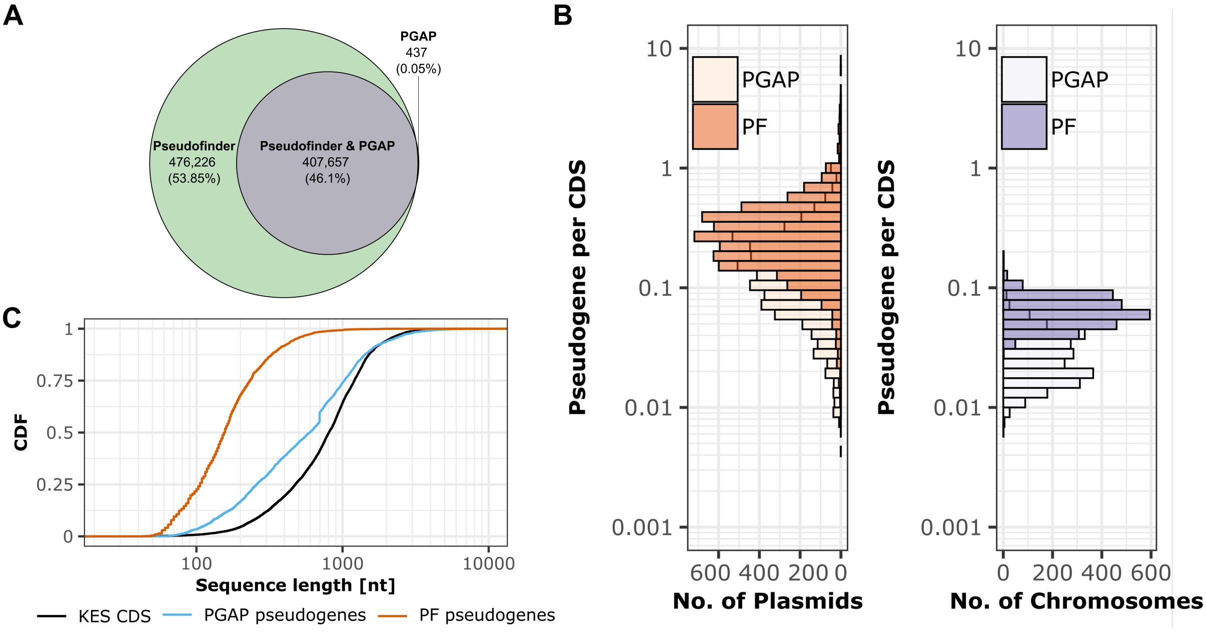

**Supplementary Fig. S1. Comparison of PGAP and Pseudofinder detected pseudogenes. (A)** Venn-Diagram of pseudogene loci determined by PGAP and Pseudofinder. Note that 437 (0.1%) PGAP pseudogene loci putatively only identified by PGAP are included in Pseudofinder's output. Pseudofinder sometimes includes two pseudogenes in one annotated locus as this tool adds a putative fragmented CDS (derived from Pseudofinder) as pseudogene downstream and upstream to annotated features (e.g., annotated PGAP pseudogenes). **(B)** Distribution of pseudogene density (pseudogene per CDS) per plasmid and chromosome for PGAP and Pseudofinder (PF) pseudogenes of the KES dataset. The graph includes plasmids and chromosomes with pseudogene content (PGAP: 5,573 (82%) plasmids and 2,442 (100%) chromosomes; Pseudofinder: 6,060 (89%) plasmids and 2,442 (100%) chromosomes). **(C)** Cumulative distribution function (CDF) of nucleotide sequence length for coding sequences, PGAP pseudogenes, and Pseudofinder pseudogenes identified in the KES dataset. Note that in **(B, C)** pseudogenes labeled as identified by PGAP have also been determined by Pseudofinder, while Pseudofinder pseudogenes have only been identified by this tool.

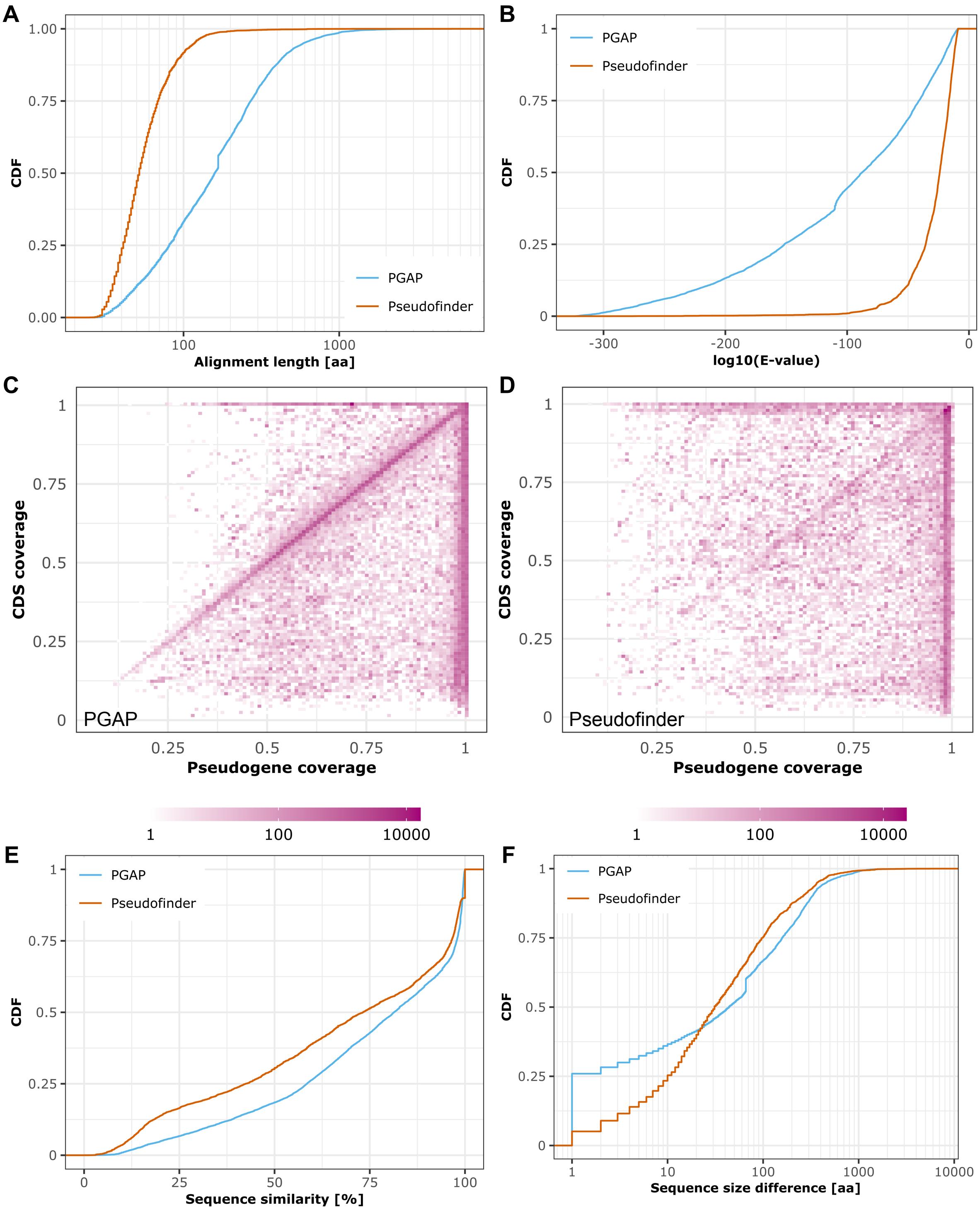

**Supplementary Fig. S2. Metrics of Pseudofinder and PGAP pseudogenes aligned to their closest homologous protein sequence. (A, B)** Cumulative distribution function (CDF) of alignment length in amino acids and E-value for local alignments of PGAP and Pseudofinder pseudogenes to their closest homologous protein sequence. **(C, D)** 2-D histograms of proportional pseudogene (x-axis) and protein sequence coverage (y-axis) for PGAP **(C)** and Pseudofinder **(D)** pseudogene local alignments to their closest homologous protein sequence. **(E, F)** CDF of sequence similarity (% of identical amino acids) and sequence size difference for global alignments of PGAP and Pseudofinder pseudogenes to their closest homologous gene sequence (in number of amino acids). Note that in **(A-F)** pseudogenes labeled as identified by PGAP have also been determined by Pseudofinder, while Pseudofinder pseudogenes have only been identified by this tool.

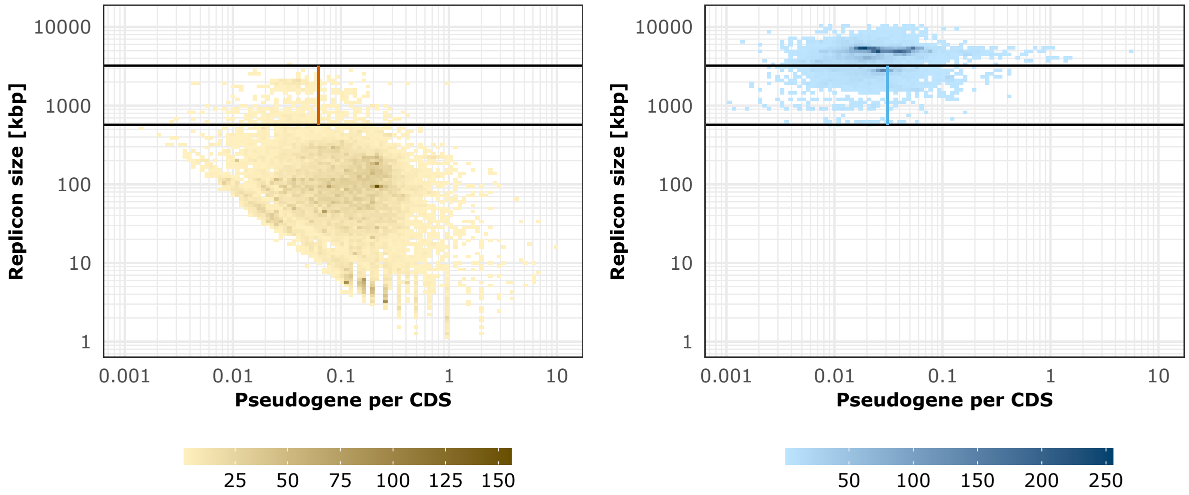

**Supplementary Fig. S3. Pseudogene density in the context of chromosomal and plasmid genome size.** The 2D-histogram plots depict pseudogene per CDS (x-axis) and replicon size (y-axis) of plasmids (left) and chromosomes (right) in the RefSeq dataset. 21,564 (74 %) plasmids and 10,828 (100 %) chromosomes with pseudogene content of the RefSeq dataset are shown. The horizontal black lines delimit a range where plasmids and chromosomes have been sampled from with a similar size distribution (tested using Kolmogorov-Smirnov, P > 0.05). Sample datasets comprising plasmids and chromosome of similar size distribution have been used for a permutation test. The vertical lines in orange (plasmids) and blue (chromosomes) represent the median ratio of pseudogene per CDS.

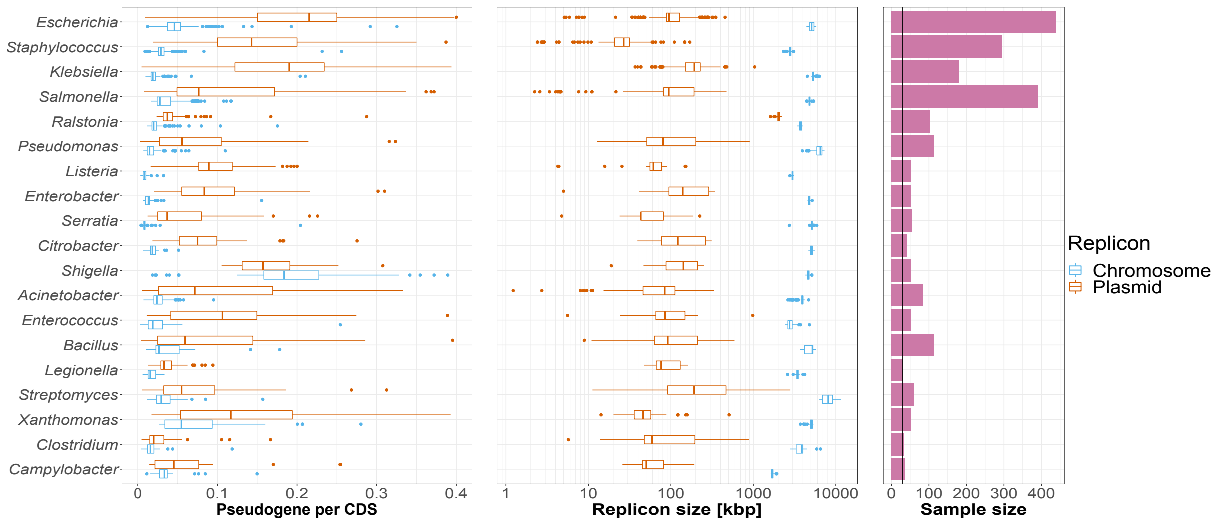

**Supplementary Fig. S4. Pseudogene density across prokaryotic genera.** Boxplots depict pseudogene per CDS and replicon size of plasmids (orange) and chromosomes (blue) per genus of the RefSeq dataset. The bars represent the sample size per genus (n>30, vertical black line). The data shown here comprises a subset of 2,238 (21 %) prokaryotic isolates from the RefSeq dataset that harbored a single chromosome and a single plasmid.

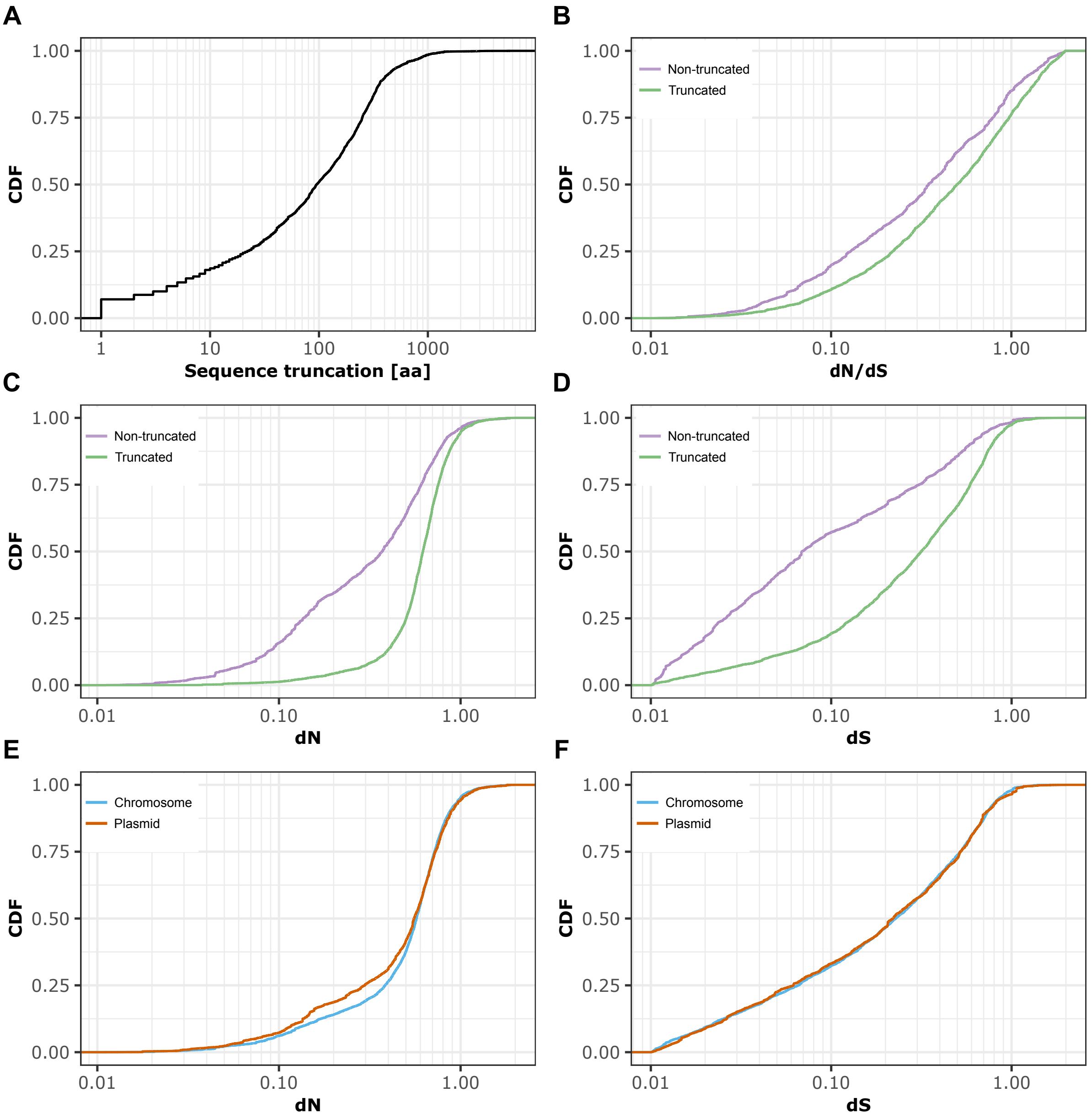

**Supplementary Fig. S5. Synonymous, and non-synonymous substitution rates of pseudogenes. (A)** Cumulative distribution function (CDF) of pseudogene sequence truncation. The pseudogene sequence truncation of the KES dataset was measured by calculating the leading and trailing gaps of the aligned translated pseudogene to the best matching homologous amino acid sequence (see also methods). 198,111 non-truncated pseudogene sequences are discarded for visual presentation. **(B, C, D)** CDF of dN/dS, dN and dS distances for truncated and non-truncated pseudogenes of the KES dataset. **(E, F)** CDF of dN and dS distances for chromosomal and plasmid pseudogenes. 110,534 pseudogenes have been removed for plotting dN, dS, and dN/dS (dS>0.01, dS<2, dN>0.01, dN<2).

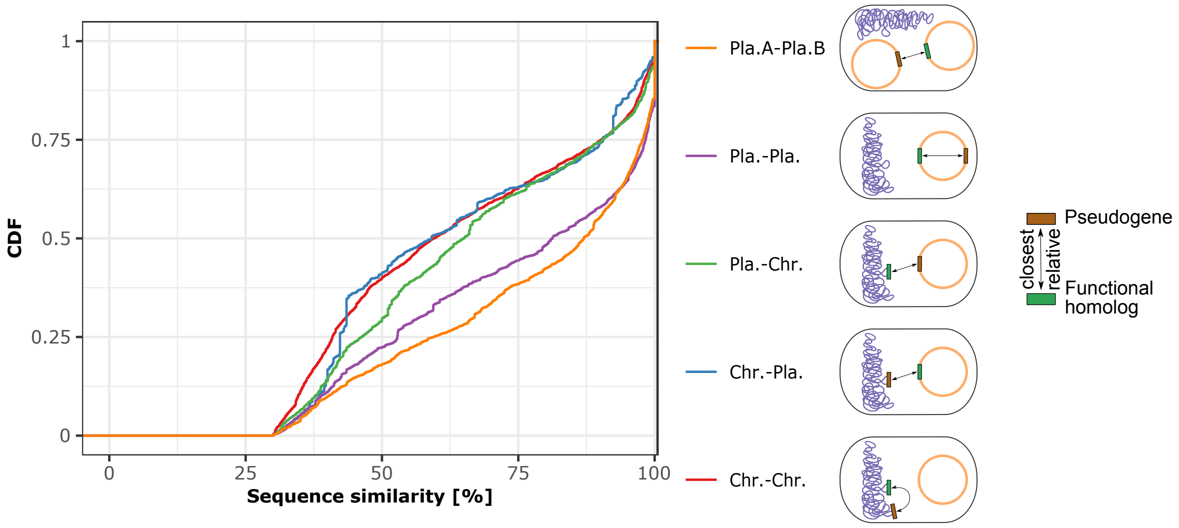

**Supplementary Fig. S6. Distribution of sequence similarity between pseudogenes and their closest homologous gene.** Cumulative distribution function (CDF) of sequence similarity (% identical amino acids) for location categories of pseudogenes to their closest homologous gene (see Fig. 1B). The location categories are sorted according to their median in descending order from top to bottom ($median_{Pla.A-Pla.B}$=87%, $median_{Pla.-Pla.}$=81%, $median_{Pla.-Chr.}$=65%, $median_{Chr.-Chr.}$=60%, $median_{Chr.-Pla.}$=59%).

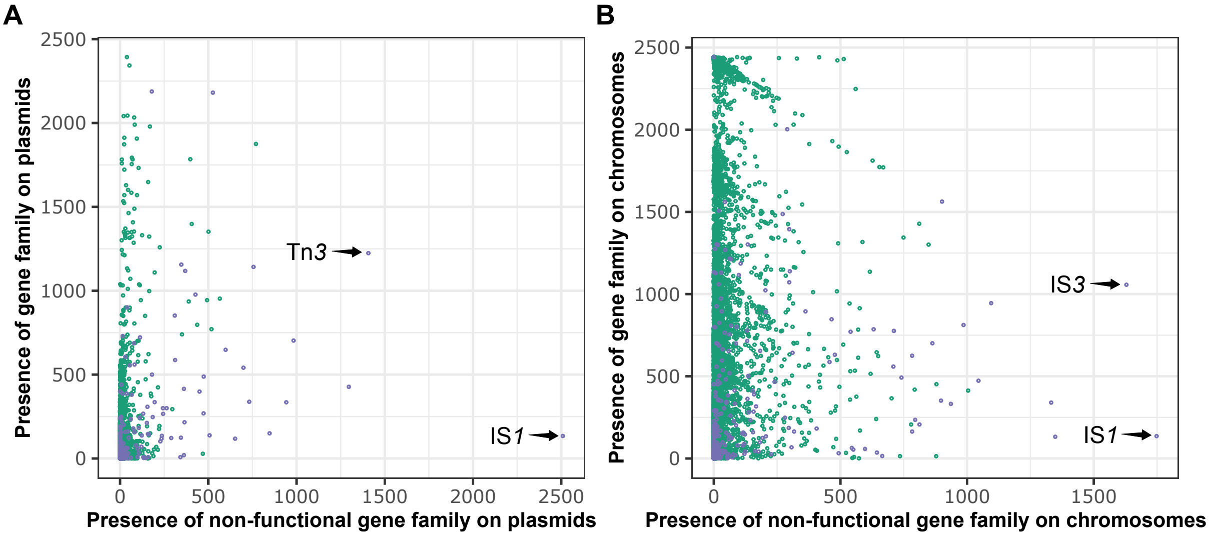

**Supplementary Fig. S7. The occurrence of gene families as coding sequences and pseudogenes. (A, B)** Presence of gene families as pseudogene (x-axis) or coding gene (y-axis) for plasmids **(A)** and chromosomes **(B)** included in the KES dataset. Transposition-related functions are overlayed and colored purple.

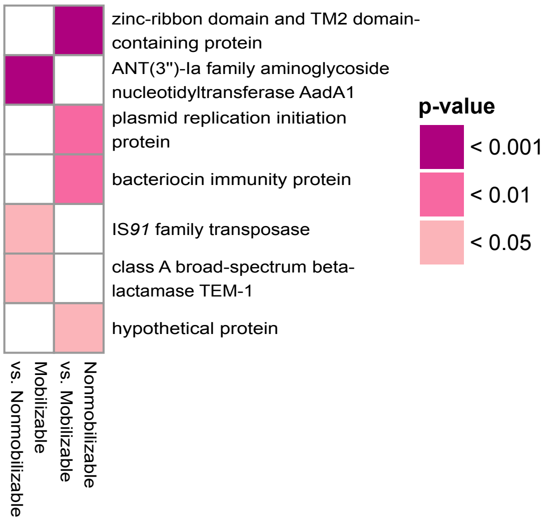

**Supplementary Fig. S8. Gene families enriched for pseudogenes in small plasmid types (<19Kb).** Non-functionalization events significantly depending on the plasmid mobility types of small plasmids (P < 0.05, using one-sided Fisher's exact test with FDR correction). First mentioned plasmid mobility type indicates in which type the enrichment of pseudogenes for gene functions has been statistically observed. The second mentioned plasmid mobility type indicates the type from which the frequencies of pseudogenes and CDSs were used for the statistical comparison.

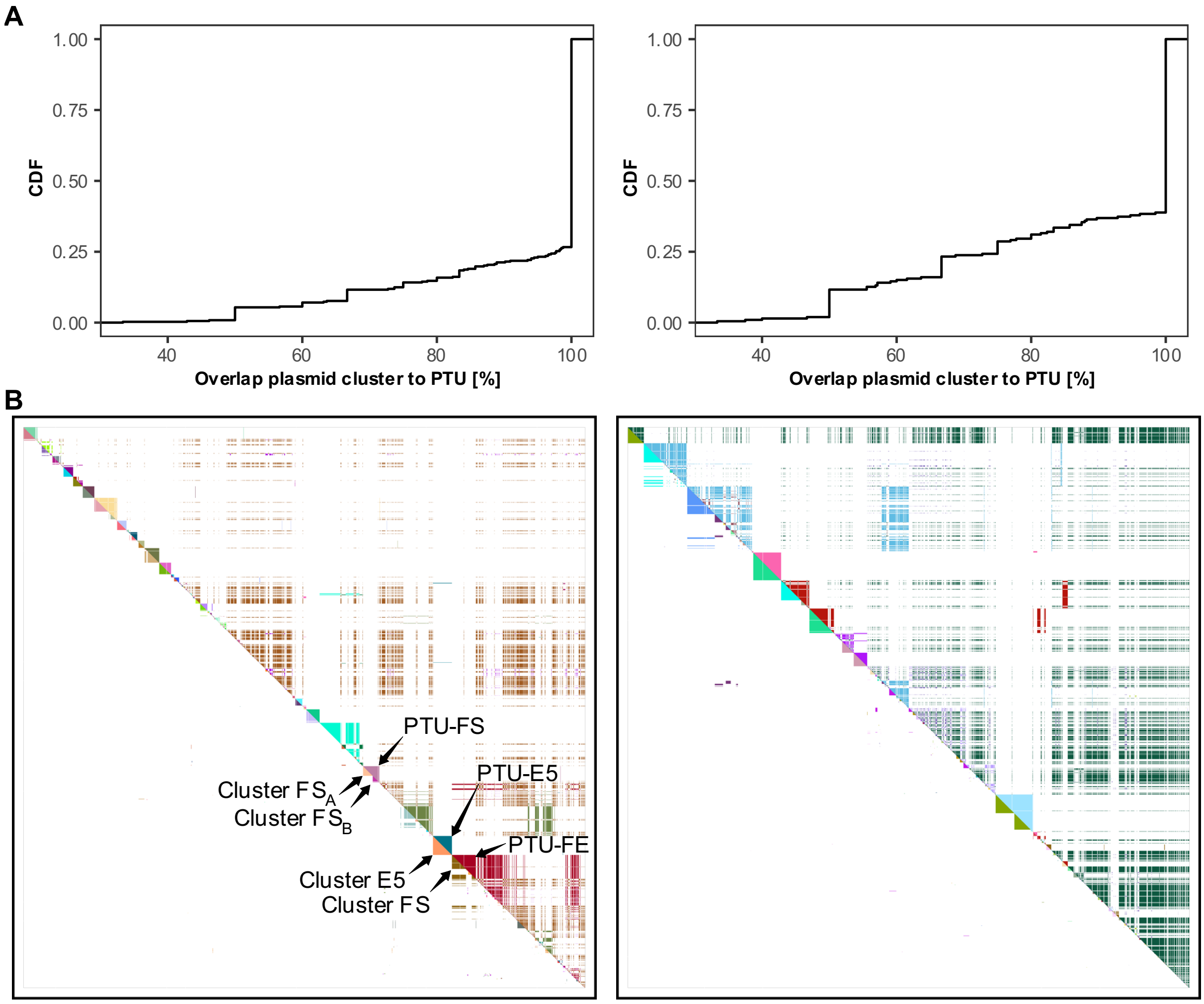

**Supplementary Fig. S9. Comparison of homologous plasmid clusters and PTUs. (A)** Cumulative distribution function of percentages a plasmid cluster matches a plasmid taxonomic unit (PTU) for large plasmid (left) and small plasmid types (right). **(B)** Plasmid clusters (lower diagonal) and PTUs (upper diagonal) depicted by distinct colors in the same order and hierarchical clustering as shown for shared gene content matrices of Fig. 3AB. The distribution of distinct PTUs in the upper diagonal indicates that some PTUs include rather distantly related plasmids.

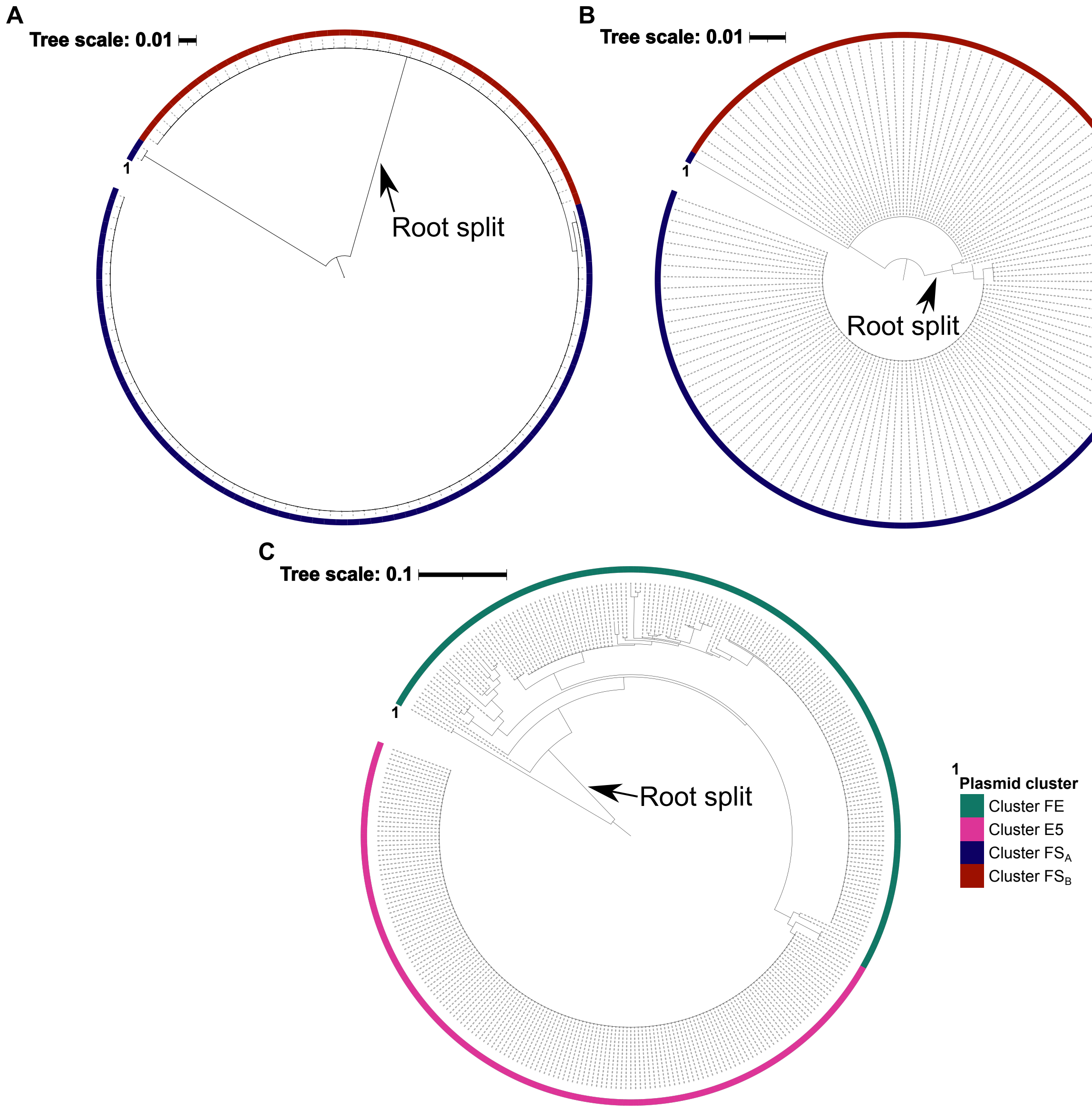

**Supplementary Fig. S10. Trees of complete single-copy genes of (homologous) plasmid clusters. (A, B, C)** Gene trees of plasmid cluster FS$_A$/FS$_B$ **(A, B)** and E5/FE **(C)** showing support of the resulting root split. The arrows indicate the root split. Outer color codes of the circular gene phylogenies depict the plasmid cluster afilitation.

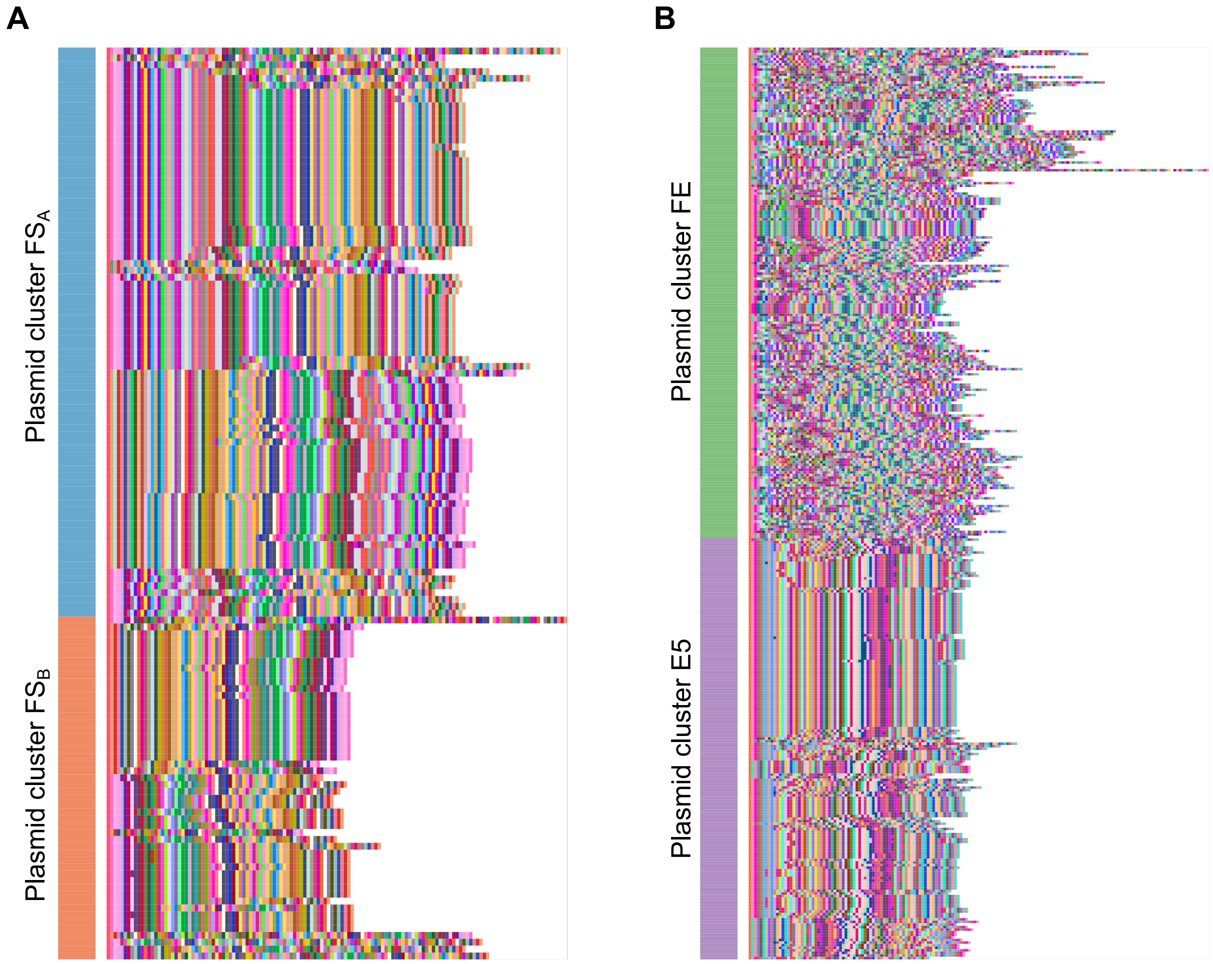

**Supplementary Fig. S11. Plasmid genome synteny of plasmid clusters. (A, B)** Synteny plots depicting conserved gene order and rearrangements in the plasmid genomes. Each row corresponds to a single plasmid genome where gene families are shown as colored rectangles along the plasmid according to their order (regardless of the gene length). The annotation bar labels corresponding plasmid clusters of the rows. The rows are sorted using hierarchically clustering according to the gene order. Universal marker gene families were used as a starting point for the plasmid plot.

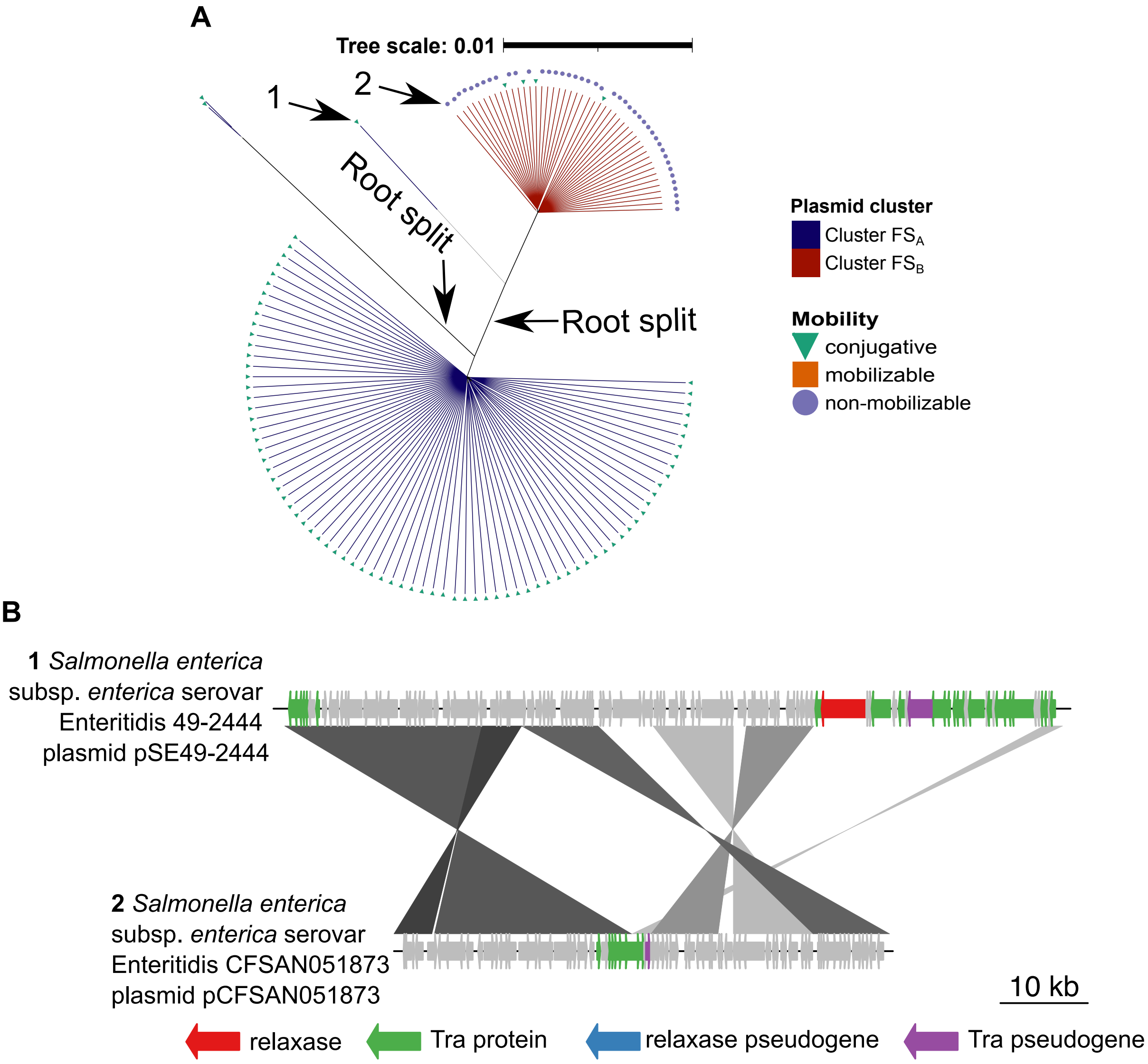

**Supplementary Fig. S12. Inference of segmental deletion of transfer-related genes in the evolution of plasmid cluster $FS_A$ and $FS_B$. (A)** Maximum-likelihood tree of plasmid cluster $FS_A$ and $FS_B$ (PTU-FS) shown in Fig. 4A. Arrows with root split labels represent the root neighborhood inference result of phylogenomic rooting. Arrows with serial numbers (1: NZ_CP018634.1, 2: NZ_CP022004.1) depict plasmids compared in **(B)**. **(B)** Blastn comparison between selected plasmids of plasmid cluster $FS_A$ and $FS_B$ (E-value $\leq 1 \times 10^{-10}$, alignment length ≥ 300bp, sequence identity ≥ 80%). Plots have been created using genoPlotR (3) (v.0.8.11). A similar observation has been made for other closely related plasmids (4).

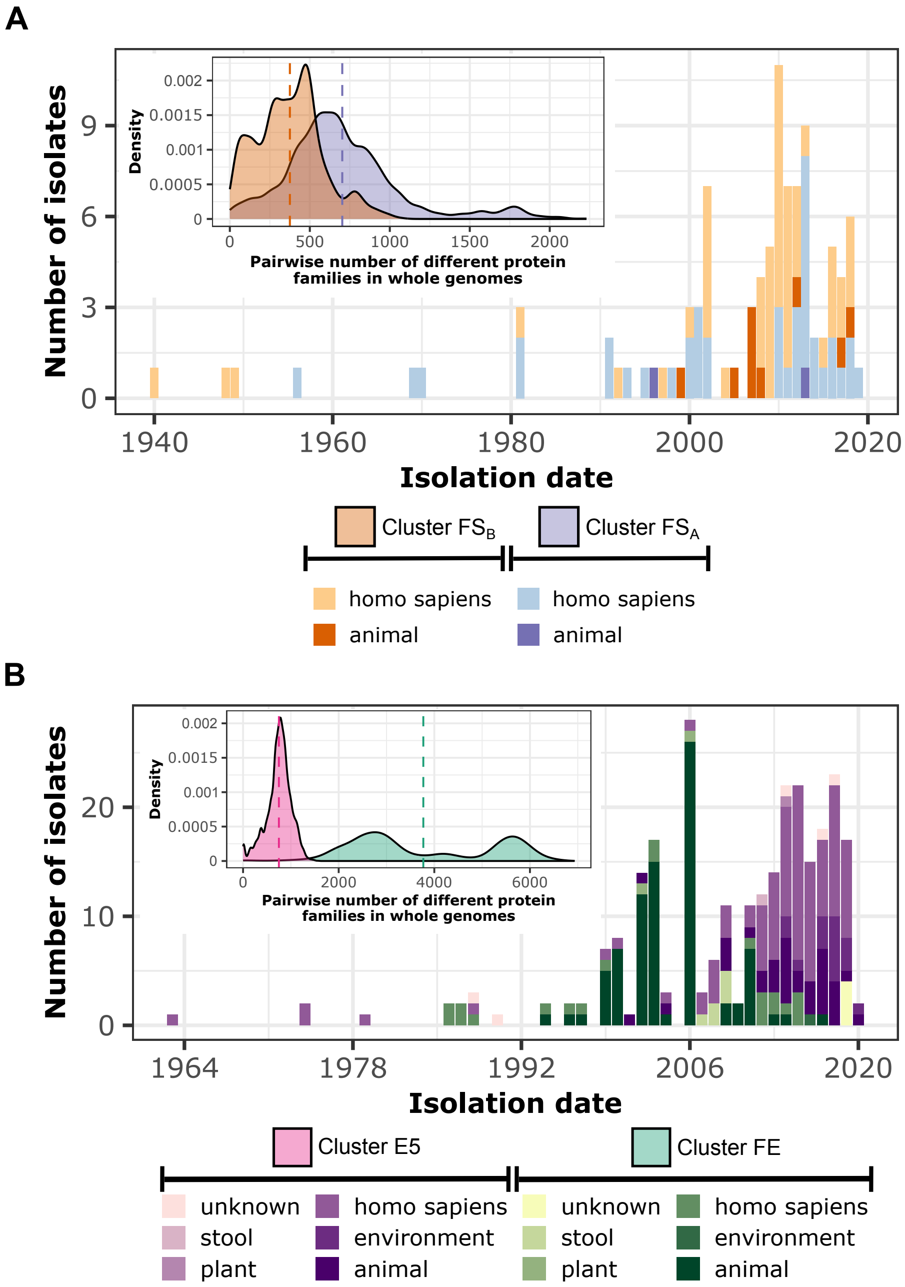

**Supplementary Fig. S13. Isolation date and divergence of hosts plasmid clusters reside in. (A, B)** Histograms show the distribution of isolation date for isolates harboring plasmid clusters $FS_A$/$FS_B$ **(A)** and FE/E5 **(B)**. The inlay density plots depict divergence measured in pairwise number of different protein families of hosts in which plasmid clusters $FS_A$/$FS_B$ **(A)** and FE/E5 **(B)** reside in. Colored dashed lines in the inlay plot represent the median of pairwise different protein families per whole genome harboring the plasmid clusters.

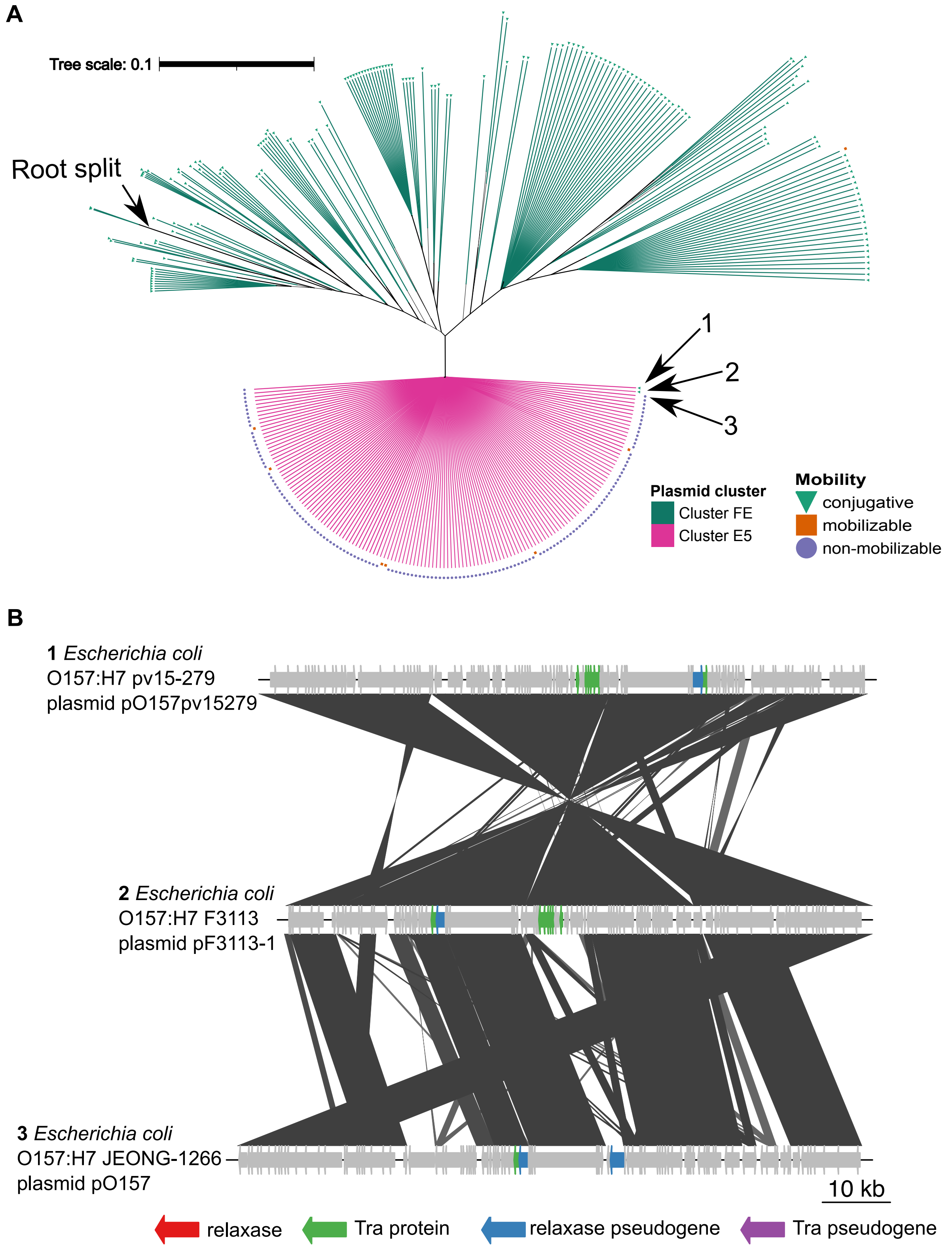

**Supplementary Fig. S14. Inference of segmental deletion of transfer-related genes in the evolution of plasmid cluster FE and E5. (A)** Maximum-likelihood tree of plasmid cluster FE (PTU-FE) and E5 (PTU-E5) shown in Fig. 4B. Arrows with root split labels represent the root inference of phylogenomic rooting. Arrows with serial numbers (1: NZ_AP018489.1, 2: NZ_CP038375.1, 3: NZ_CP015816.1) depict plasmids compared in **(B)**. **(B)** Blastn comparison between selected plasmids of plasmid cluster FE and E5 (E-value ≤ $1 \times 10^{-10}$, alignment length ≥ 300bp, sequence identity ≥ 80%). Plots have been created using genoPlotR (3) (v.0.8.11).
